# The UBR-1 Ubiquitin Ligase Regulates Glutamate Metabolism to Generate Coordinated Motor Pattern in *C. elegans*

**DOI:** 10.1101/198994

**Authors:** Jyothsna Chitturi, Wesley Hung, Anas M. Abdel Rahman, Min Wu, Maria A. Lim, John Calarco, Renee Baran, Xun Huang, James W. Dennis, Mei Zhen

## Abstract

UBR1 is an E3 ubiquitin ligase best known for its ability to target protein degradation by the N-end rule. The physiological functions of UBR family proteins, however, remain not fully understood. We found that the functional loss of *C. elegans* UBR-1 leads to a specific motor deficit: when adult animals generate reversal movements, A-class motor neurons exhibit synchronized activation, preventing body bending. This motor deficit is rescued by removing GOT-1, a transaminase that converts aspartate to glutamate. Both UBR-1 and GOT-1 are expressed and critically required in premotor interneurons of the reversal motor circuit to regulate the motor pattern. *ubr-1* and *got-1* mutants exhibit elevated and decreased glutamate level, respectively. These results raise an intriguing possibility that UBR proteins regulate glutamate metabolism, which is critical for neuronal development and signaling.

**Author Summary:** Ubiquitin-mediated protein degradation is central to diverse biological processes. The selection of substrates for degradation is carried out by the E3 ubiquitin ligases, which target specific groups of proteins for ubiquitination. The human genome encodes hundreds of E3 ligases; many exhibit sequence conservation across animal species, including one such ligase called UBR1. Patients carrying mutations in *UBR1* exhibit severe systemic defects, but the biology behinds UBR1’s physiological function remains elusive. Here we found that the *C. elegans* UBR-1 regulates glutamate level. When UBR-1 is defective, *C. elegans* exhibits increased glutamate; this leads to synchronization of motor neuron activity, hence defective locomotion when animals reach adulthood. UBR1-mediated glutamate metabolism may contribute to the physiological defects of *UBR1* mutations.

## Introduction

In eukaryotic cells, the ubiquitin-proteasome system mediates selective protein degradation (1, 2). E3 ubiquitin ligases confer substrate specificity via selective interaction with the degradation signals in substrates (3-6).

UBR1 acts not only as an E3 ligase for the N-end rule substrates, whose metabolic stability is determined by the identity and post-translational status of their N-terminal moiety, but also for substrates that do not harbor the N-terminal degrons (6, 7). The UBR family proteins exist from yeast to man, and have been implicated in multiple cellular processes (reviewed in 7). Yeast UBR1 is not essential, but *ubr1* mutants exhibit less efficient chromatin separation, and mildly increased doubling time (8). Yeast UBR1 also participates in protein quality control, potentiating the degradation of mis-folded proteins by ER membrane ligases (9). The loss of *C. elegans* UBR-1 results in delayed degradation of a regulator for post-embryonic hypodermic cell division, but does not cause obvious hypodermic defect (10). In mammalian cell lines, the N-end rule pathway targets pro-apoptosis fragments for degradation, affecting the efficacy of induced apoptosis (11). The simultaneous loss of two rodent UBR homologues results in embryonic lethality with severe developmental defects in the heart and brain (12). In human, loss-of-function mutations in one of several UBR family proteins, UBR1, cause the Johanson-Blizzard Syndrome (*JBS*), a genetic disorder with multi-systemic symptoms including pancreatic insufficiency, growth retardation, and cognitive impairments (13). To date, a unifying physiological function of UBR proteins in animal models and human is lacking. In fact, whether UBR1’s role as an N-end rule E3 ligase is relevant for the *JBS* pathophysiology remains elusive (14).

*C. elegans* has a single UBR1 ortholog, UBR-1. Using this simplified animal model, we reveal that the functional loss of UBR-1 leads to a specific, late onset and prominent motor pattern change, and such a change is reversed by removing a metabolic enzyme, GOT-1, which we find to synthesize glutamate from aspartate.

Glutamate is an abundant amino acid. As a metabolite, it is essential for metabolism and development. As a neurotransmitter, glutamate-mediated signaling regulates animals’ motor and cognitive functions (15-18). As such, glutamate level needs to be tightly regulated for metabolism, as well as neuronal signaling. Aberrant glutamate signaling has been implicated in cytotoxicity (19), and neurological disorders (20).

The genetic interaction between *ubr-1* and *got-1* mutants suggests that UBR-1 may affect glutamate metabolism. Consistent with this notion, we find that both genes are critically required in premotor interneurons to effect the reversal motor pattern change. Further, our metabolomics analyses reveal an inversely correlated change - increased and decreased glutamate level - in *ubr-1* and *got-1* mutant animals, respectively. These findings reveal a previously unknown role for the UBR family protein in glutamate metabolism. They further allude to the possibility that a common cellular defect, such as that in glutamate metabolism, may contribute to UBR’s multi-systemic functions.

## Results

### UBR-1 promotes bending during reversal movements

Wildtype *C. elegans* generates movements through propagating body bends. In a genetic screen for mutants with altered motor patterns, we isolated *hp684* (Fig. 1A), a mutant that is capable of reversal movements, but does so with limited body flexing (Fig. 1B; Supplementary Movie 1). The stiffness is prominent during prolonged reversals, and is progressive as animals develop from the last-stage larvae into adults.

**Figure 1.**
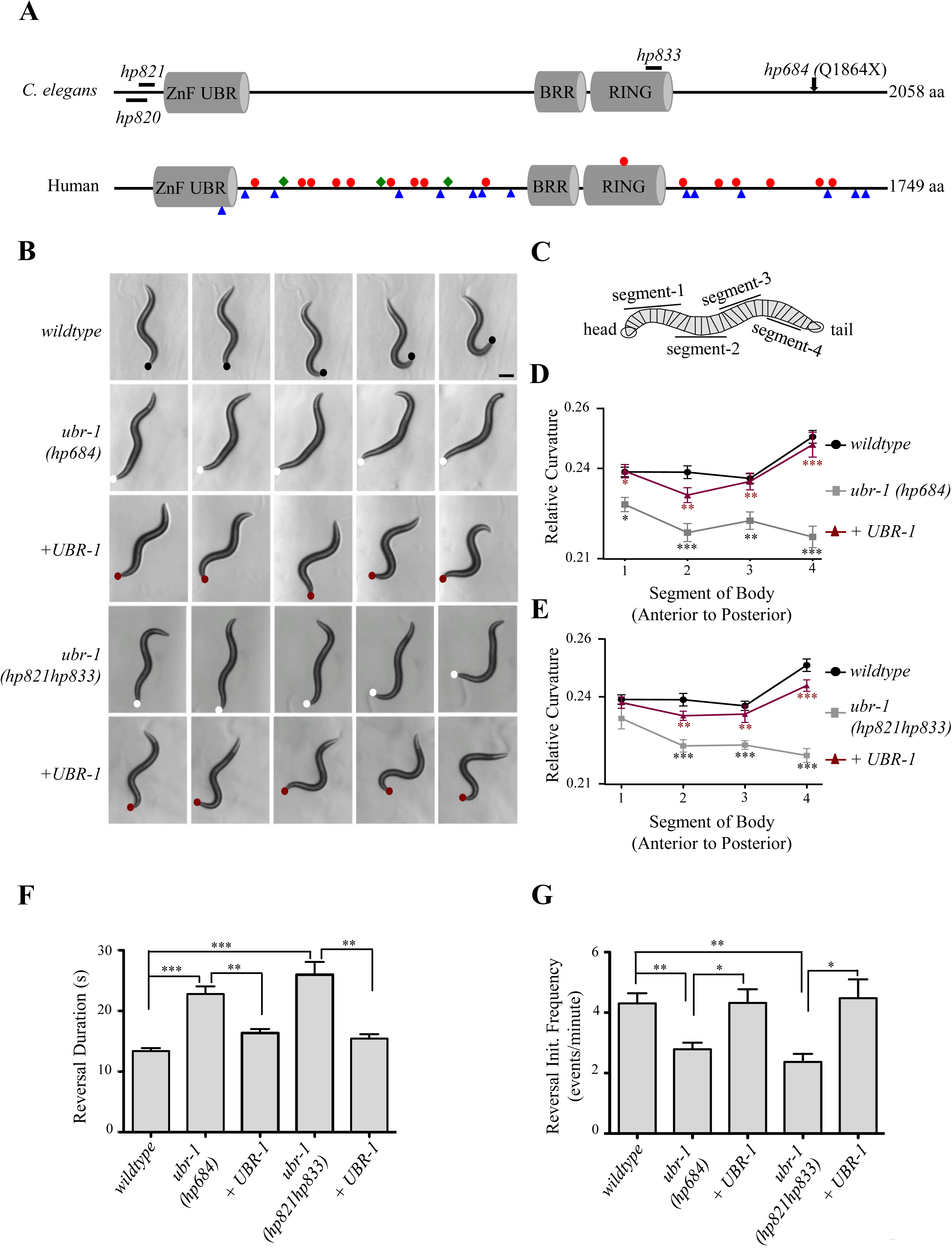
*ubr-1* mutants exhibit reduced bending during reversal locomotion. A) Structure of the UBR1 family protein. Position and amino acid substitutions in *C. elegans* alleles are denoted in the upper panel. Nonsense mutations (red dots), frame shift mutations (blue triangles), and small in-frame deletions (green diamonds) in human UBR1 that lead to *JBS* are denoted in the lower panel. B) Consecutive images of a wild type and *ubr-1* animal during reversals (left to right panels). Wildtype animals generate sinusoidal body bends, whereas *ubr-1(hp684, hp821hp833)* animals exhibit almost no bending, which was rescued by restoring expression of UBR-1. Dots denote position of tail. Scale bar 200µm. C) Schematic of curvature analysis. A worm skeleton is divided into 29 segments, and relative curvature is calculated for each segment from anterior to posterior, and binned into 4 groups. D, E) In *ubr-1* mutants (grey line), bending curvature is reduced across all segments compared to wildtype (black line), and this is rescued by restoring UBR-1 expression (red line). D-*ubr-1(hp684)*; E-*ubr-1(hp821hp833)*. *ubr-1* mutants exhibited longer (F) reversal events, and reduced reversal initiation frequency (events/per minute) (G). *P<0.05, ** P<0.01, ***P<0.001 by Two-way RM ANOVA (D, E), and by the Kruskal-Wallis test (F, G). Data are represented as mean ± SEM.

We quantified the motor phenotypes of one-day-old wildtype and *hp684* adults. Because the forward movement is the preferred motor state under laboratory conditions, we assayed their motor behaviors on plates where they were frequently induced for prolonged reversals (Methods). We compared their body curvature and duration of reversals during these events (Fig. 1D-F). *hp684* mutants exhibited significantly decreased mean curvature during reversals (Fig. 1D), significantly longer duration (Fig. 1F), and reduced reversal initiation frequency (Fig. 1G) under the assay conditions (Methods). The bending curvature and duration of forward movements, captured during the same recording periods, were only mildly affected or unchanged in *hp684* mutant animals (Supplementary Movie 1).

We mapped and identified the causative genetic lesion in *hp684* mutants (Methods): a recessive and nonsense mutation that leads to truncation of the last 194 amino acids (Q1864X) of UBR-1 (Fig. 1A). The UBR family proteins exhibit conserved domain organization from yeast to humans. In addition to a highly conserved C-terminal sequences, they share the N-terminal UBR box, and internal motifs that include a region enriched for basic amino acids (BRR), and a RING finger (21). The UBR box and its neighboring sequences interact with the substrates and E2 ubiquitin-conjugating enzyme of the N-end rule (22), whereas the RING finger is the hallmark motif utilized by a large class of E3 ligases to recruit non-N-end rule substrates (23-25).

To further verify that the motor phenotypes that we observed in *hp684* mutants result from the functional loss of UBR-1, we generated multiple *ubr-1* deletion alleles, *hp820, hp821*, and *hp821hp833* (Fig. 1A), by CRISPR-Cas9-mediated genome editing (26) (Methods). *hp820* harbors a small in-frame deletion near the N-terminus (ΔR18-W25); *hp821* a N-terminal four base pair deletion that leads to a frame-shift and a premature, N-terminal stop codon (E34X), and *hp821hp833*, in addition to *hp821*, a seven-base-pair deletion in the RING finger that leads to a frame shift and premature internal stop codon (E1315X). All alleles, like *hp684*, are recessive and viable, all exhibited motor defects similar to *hp684,* including reduced bending, increased duration, and reduced initiation frequency during reversal movements (Fig. 1B, 1E-G; Fig. S1). Also like *hp684*, their reversal defects were rescued by a genomic fragment that harbors only *ubr-1* (Fig. 1B-G; Fig. S1).

These results confirm that the motor defects exhibited by all *ubr-1* alleles result from the functional loss of UBR-1. The comparable phenotypic severity among *hp684, hp820, hp821* and *hp821hp833* mutants complements findings from the *JBS* patients, where diverse UBR1 mutations, resulting in a wide range of genetic lesions - early or late stop codons, reading frame shifts, and small internal in-frame deletions (Fig. 1A) - exert similar pathologic effects (27).

Hereafter, we present quantified results from the *hp684* allele unless specified, and refer to it as the *ubr-1* mutants. For behavioral analyses, we present data for reversal movements, and refer to it as the motor phenotype.

### A critical requirement of UBR-1 in premotor interneurons to promote bending

To begin probing UBR-1’s function, we first examined the expression pattern of a GFP::UBR-1 reporter, which fully reversed *ubr-1* mutants’ motor defects. GFP::UBR-1 exhibits strong expression in a fraction of neurons and all musculatures throughout development, and weak expression in hypodermal seam cells (Fig. 2A). In the motor circuit, UBR-1::GFP’s expression is prominent in premotor interneurons (INs in Fig. 2A) of the reversal motor circuit (Fig. 2A; Fig. S2A), and is absent from all ventral cord motor neurons that execute locomotion.

**Figure 2.**
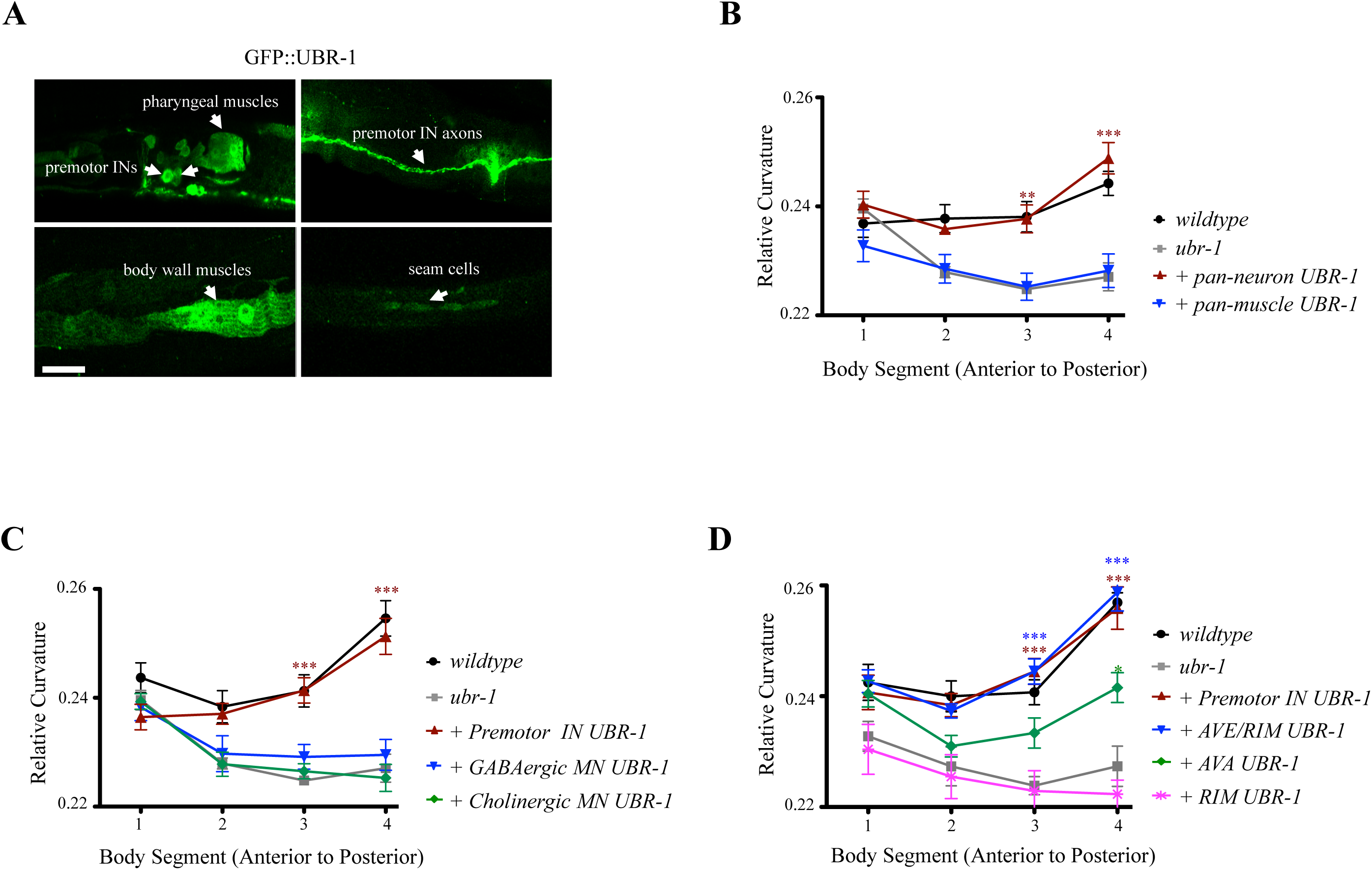
A critical requirement of UBR-1 in premotor interneurons during reversals. A) The UBR-1::GFP reporter was expressed in a subset of neurons including the premotor interneurons of the reversal circuit (Premotor INs), the pharyngeal muscles, the body wall muscles, and hypodermal seam cells. Scale bar 10µm. B) Reduced bending in *ubr-1* mutants (grey line) was rescued by restoring UBR-1 expression in neurons (red line), but not in muscles (blue line). C) Reduced bending in *ubr-1* mutants (grey line) was robustly rescued by restoring UBR-1 expression in glutamate-receptor-expressing, premotor interneurons (INs), not in cholinergic or GABAergic motor neurons (MNs). D) Among the premotor interneurons (INs), a robust rescue required UBR-1 expression AVE and RIM (red line). *P<0.05, ** P<0.01, ***P<0.001 by Two-way RM ANOVA. Data are represented as mean ± SEM.

To determine the critical cellular origins of *ubr-1* mutants’ motor defect, we examined the effect of restoring UBR-1 using exogenous and cell-type specific promoters. When UBR-1 expression was restored panneuronally (*Prgef-1*), the reversal motor defects (Fig. 2B) in *ubr-1* mutants were fully rescued, whereas restoring UBR-1 in muscles (*Pmyo-3*) did not (Fig. 2B; Fig. S3A, B). These results show a neuronal origin of *ubr-1* mutants’ motor phenotype.

Through examining the effect of restoring UBR-1 expression in subgroups of motor circuit neurons that partially overlap with identified UBR-1::GFP-positive neurons, we confirmed a critical requirement of UBR-1 in premotor interneurons (Fig. 2C; Fig. S3A, B). Specifically, restoring UBR-1 expression by either *Pglr-1* or *Pnmr-1* significantly rescued *ubr-1* mutants’ motor defects, including bending, duration, and frequency during reversal movements (Fig. 2C; Fig. S3A, B), whereas restoring UBR-1 in motor neurons did not (Fig. 2C; Fig. S3A, B). Both *Pglr-*1 *and Pnmr-1* activate expression in premotor interneurons of the reversal motor circuit, AVA, AVE, AVD, and RIM (28-30). Their activation, inactivation, and ablation affect the execution and characteristics of the reversal motor states (31-43). Importantly, restoration of UBR-1 in other interneurons, including the premotor interneurons of the forward motor circuit (AVB), did not rescue *ubr-1* mutants’ motor phenotype (Table 1).

**Table 1.** **The effect of restored expression of UBR-1 and GOT-1 on *ubr-1* and *ubr-1; got-1*’s reversal motor pattern.**

| Background | Constructs (transgenes) | Tissues with restored expression | Reversals |
| --- | --- | --- | --- |
| *ubr-1* | - | - | - |
|  | *Pubr-1::UBR-1* | neurons, muscles, seam cells | +++ |
|  | *Prgef-1::UBR-1* | all neurons | +++ |
|  | *Pmyo-3::UBR-1* | Muscles | - |
|  | *Pglr-1::UBR-1* | premotor INs (AVA, **AVE**, **RIM**, AVD, PVC, AVB); others | +++ |
|  | *Pnmr-1::UBR-1* | premotor INs (AVA, **AVE**, **RIM**, AVD, PVC); others | +++ |
|  | *Popt-3::UBR-1* | premotor INs (**AVE**, **RIM**); others | +++ |
|  | *Prig-3::UBR-1* | premotor INs (AVA); others | + |
|  | *Pgcy-13::UBR-1* | premotor INs (RIM) | - |
|  | *Punc-47::UBR-1* | GABAergic MNs | - |
|  | *Pacr-2::UBR-1* | cholinergic MNs; others | - |
|  | *Plgc-55::UBR-1* | premotor INs (AVB); others | - |
|  | *Pttx-3::UBR-1* | INs (AIY) | - |
|  | *Pinx-1::UBR-1* | INs (AIB) | - |
| *ubr-1*;<br>*got-1* | - | - | +++ |
|  | *Prgef-1::GOT-1* | all neurons | - |
|  | *Pglr-1::GOT-1* | premotor INs (AVA, **AVE**, **RIM**, AVD, PVC, AVB); others | - |
|  | *Popt-3::GOT-1* | premotor INs (**AVE**, **RIM**); others | - |
|  | *Prig-3::GOT-1* | premotor INs (AVA); others | + |
|  | *Pgcy-13::GOT-1* | premotor INs (AVA); others | +++ |
|  | *Punc-47::GOT-1* | GABAergic MNs | +++ |

Within this group of premotor interneurons of the reversal circuit, restoration of UBR-1 in AVE and RIM (*Popt-3*) exerted the strongest partial rescue (Fig. 2D; Fig. S3C, D), whereas the restoration in AVA (*Prig-3*), or RIM (*Pgcy-13*) alone, led to modest to no rescue (Fig. 2D; Fig. S3C, D). UBR-1’s role in the reversal motor circuit involves the whole network of premotor interneurons, with AVA, AVE, and maybe RIM being the most critical components.

### UBR-1 promotes bending by preventing simultaneous A motor neuron activation

What underlies the reduced bending in *ubr-1* mutants? Premotor interneurons of the reversal motor circuit innervate the A-class motor neurons (A-MNs) (Fig. 3A). Multiple A-MNs innervate body wall muscles to execute reversal movements: they are divided into the ventral (VA) and dorsal (DA) muscle-innervating subtypes through likely non-overlapping neuromuscular junctions (32, 34, 44, 45). We examined the temporal activation pattern of a posterior cluster A-MNs, DA7, VA10, and VA11 (predicted muscle targets illustrated in Fig. 3B) in freely moving adults that express a calcium sensor GCaMP6s::wCherry (46) in these A-MNs (Methods).

**Figure 3.**
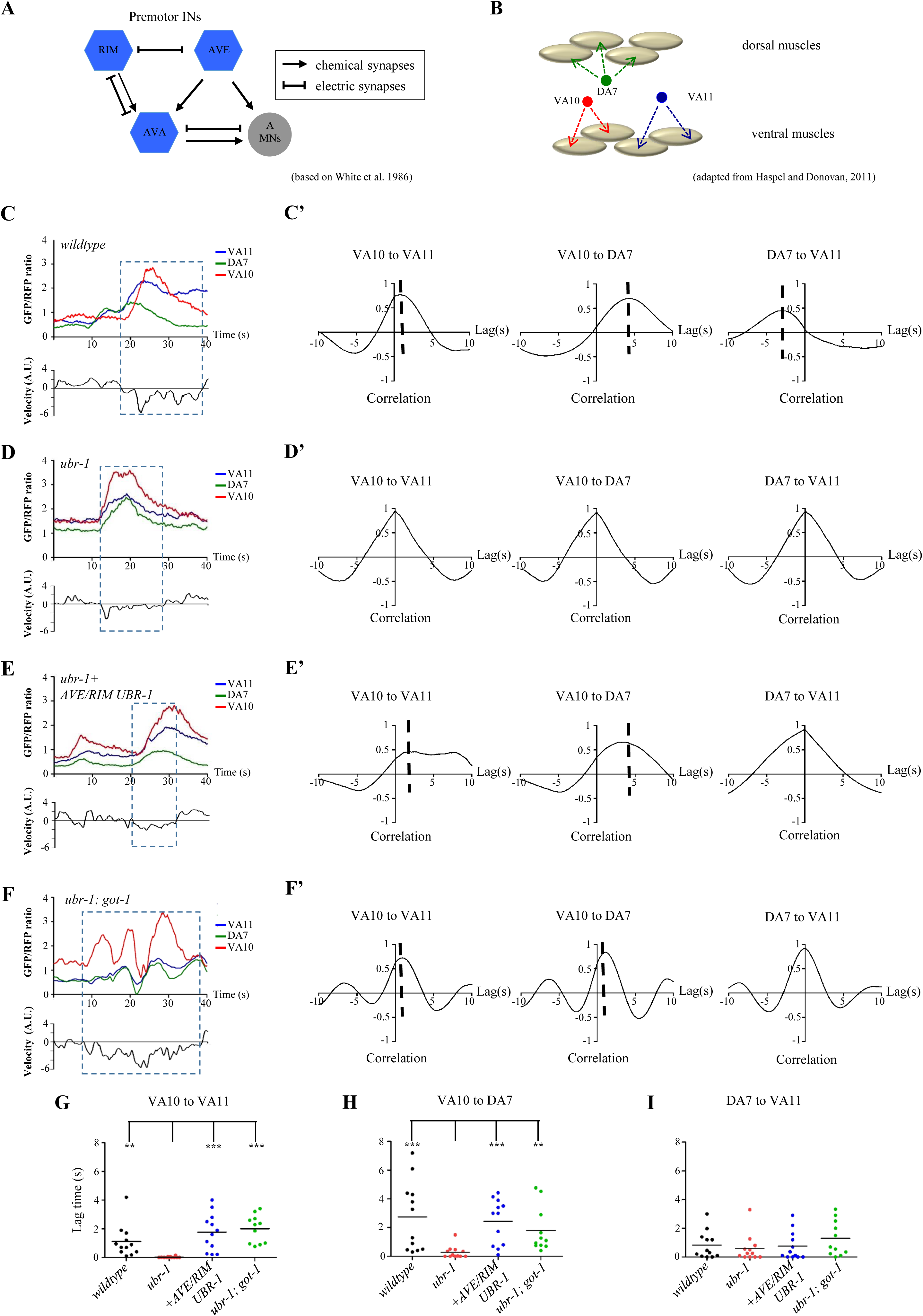
UBR-1 promotes body bending by preventing synchronized A motor neuron activation. A) Diagram of connectivity of the *C. elegans* reversal motor circuit, based on (45). Hexagons and the circle denote premotor interneurons, and the A class motor neurons, respectively. Arrows and lines denote chemical and electrical synapses between neurons, respectively. B) Approximated anatomic positions of the three A class motor neurons (VA11, DA7, VA10) and their predicted ventral and dorsal muscle targets (adapted from (79). C) An example trace of simultaneous calcium imaging of three A-type motor neurons in a moving wildtype animal. Upper panel: activities of VA11, DA7, and VA10 neurons, reflected by the GCaMP6/RFP signal ratio (Upper panel); Lower panel: the instantaneous velocity of the animal, reflected by the displacement of DA7 soma position (Lower panel; positive values indicate moving towards the head; negative values indicate moving towards the tail), during a period of 40s from a 3min recording. Boxed period denotes the reversal period applied for cross-correlation analyses. C’) The cross-correlation of the activity profiles between VA10 and VA11 (left), VA10 and DA7 (center), and DA7 and VA11 (right), respectively. Dotted vertical line denotes the lag time. D, D’) An example trace of simultaneous calcium imaging (D) and cross-correlation analysis (D’) of VA11, DA7 and VA10 in a moving *ubr-1* animal. E, E’) An example trace of simultaneous calcium imaging (E) and cross-correlation analysis (E’) of VA11, DA7 and VA10 in a moving transgenic *ubr-1* mutant animal with restored UBR-1’s expression in neurons that include AVE/RIM premotor interneurons. F, F’) An example trace of simultaneous calcium imaging (F) and cross-correlation analysis (F’) of VA11, DA7 and VA10 in a moving *ubr-1; got-1* mutant animal. G-I) The phase lags between activities of VA10 and VA11 (G), VA10 and DA7 (H), and DA7 and VA11 (I) in animals of respective genotypes. The asynchrony between VA10 and VA11 (G), and between VA10 and DA7 (H) are significantly reduced in *ubr-1* mutants compared to wildtype animals, and restored in *ubr-1* mutants by both UBR-1 expression in premotor interneurons, and the *got-1* mutation. The activation of DA7 and VA11 (I), with higher synchrony than the other two pairs in wildtype animals, was not significantly altered in *ubr-1* mutants. *P<0.05, ** P<0.01, ***P<0.001by the Kruskal-Wallis test. Horizontal lines represent mean values.

Consistent with the notion that A-MNs execute reversal movements, they exhibited calcium changes when animals moved backwards (boxed area in Fig. 3C). While the frequency and amplitude of the calcium waveforms for A-MNs varied for each reversal event (47), their calcium profiles exhibited phase relations that are consisted with the expected temporal activation of muscle groups that they are predicted to innervate. Specifically, VA10 and VA11, two A-MNs that innervate adjacent ventral muscles exhibited asynchrony in activation (red and blue traces in Fig. 3C; left panel in Fig. 3C’), with variable lags (Fig. 3G; Methods), as expected from the sequential contraction of adjacent muscles during reversals at different velocities. DA7 likely innervates dorsal muscles that appose VA10 and V11’s ventral targets (Fig. 3B). Consistent with the alternating dorsal and ventral muscle contraction during bending, VA10 and DA7 exhibited asynchrony of activity patterns (green and red traces in Fig. 3C; middle panel in Fig. 3C’) with variable lags (Fig. 3H). DA7 and VA11 were activated in relative synchrony (green and blue traces in Fig. 3C; right panel in Fig. 3C’), exhibiting shorter lags (Fig. 3I) when compared to the other A-MN pairs. This indicates a more direct dorsal-ventral opposition between VA10 and DA7’s muscle targets.

We observed a striking difference in A-MN’s activation pattern in *ubr-1* mutants. While they also exhibited calcium changes during reversals, all three A-MNs’ activation exhibited synchrony (blue, red and green traces in Fig. 3D; Fig. 3D’). This led to a drastic reduction in the mean lag times between VA10 and VA11 (Fig. 3G), and between VA10 and DA7 (Fig. 3H), whereas the short lags between DA7 and VA11 remained statistically unchanged (Fig. 3I) between wildtype and *ubr-1* mutants.

Hence, reduced bending in *ubr-1* mutants was caused by increased synchronization, not lack of A-MNs’ activities. Importantly, when UBR-1 was restored in the premotor interneurons of the reversal circuit, A-MN’s phasic relationships were restored in *ubr-1* mutants (Fig. 3E, E’, Fig. G-I). Therefore UBR-1 plays a critical role in premotor interneurons to ensure sequential motor neuron activation, which underlies bending during reversal movements.

### Removing the GOT-1 transaminase restores *ubr-1* mutants’ bending

If UBR-1, an E3 ligase, affects the animal’s motor pattern through negative regulation of a biological pathway, the pattern change should be rescued by a simultaneous decrease of the activity of the pathway. Accordingly, we screened for genetic suppressors of *ubr-1* mutants’ motor phenotype.

We isolated *hp731*, which restored bending in *ubr-1* mutants during reversal movements (Fig. 4C; Supplementary Movie 2). Notably, both *hp731* and *ubr-1; hp731* mutant animals exhibited slightly deeper bending than wildtype animals (Fig. 4C; Supplementary Movie 2). *hp731* also significantly rescued *ubr-1* mutant’s change in reversal duration and frequency (Fig. S4A, B). Consistent with synchronized A-MN activation underling *ubr-1*’s lack of bending, VA10, DA7 and VA11’s phasic relationships were restored in *ubr-1; hp731* mutants (Fig. 3F-I).

**Figure 4.**
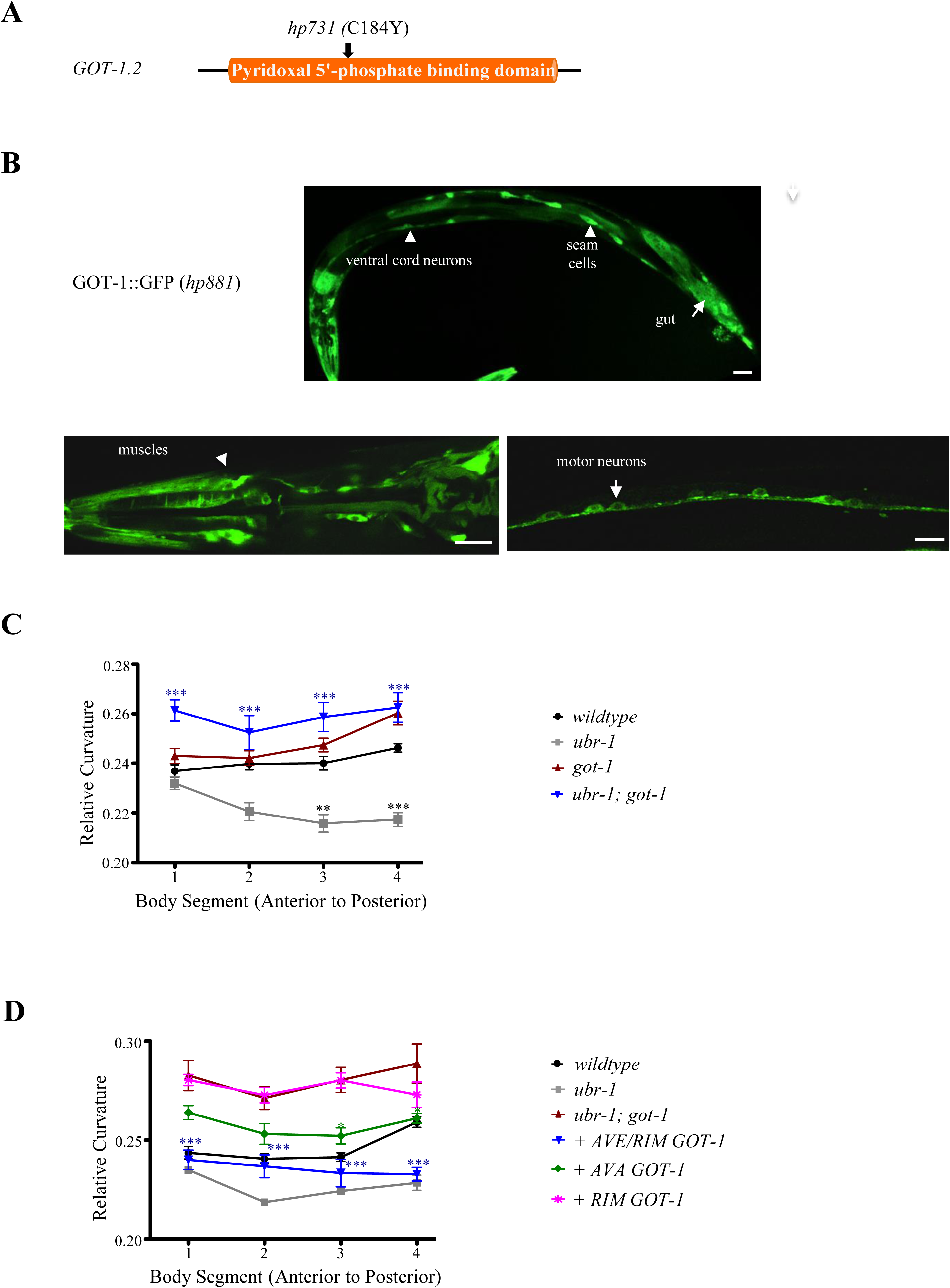
Removing GOT-1 transaminase restores bending in *ubr-1* mutants. A) A diagram depicting the structure of GOT-1 and the molecular lesion of the *hp731* allele. B) GOT-1 is ubiquitously expressed in all somatic tissues. Scale bar 10µm. C) Mutations in *got-1* suppress motor neuron defects in *ubr-1*. D) Restoring GOT-1 in premotor INs including AVE and RIM (blue line) in *ubr-1; got-1* reverted curvature to that of *ubr-1*. *P<0.05, ** P<0.01, ***P<0.001 by the Two-way RM ANOVA test. Data are represented as mean ± SEM.

We identified the causative mutation in *hp731* mutant animals. *hp731* harbors a causative, *lf,* and missense C184Y mutation (Methods) at the pyridoxal phosphate-binding domain in GOT-1.2 (Fig. 4A), one of the four predicted *C. elegans* glutamate-oxaloacetate transaminases (GOTs). GOT enzymes catalyze the transfer of an amino group between aspartate and α-ketoglutarate (α-KG) and convert them to oxaloacetate (OAA) and glutamate (48, 49) (Fig. 5A).

**Figure 5.**
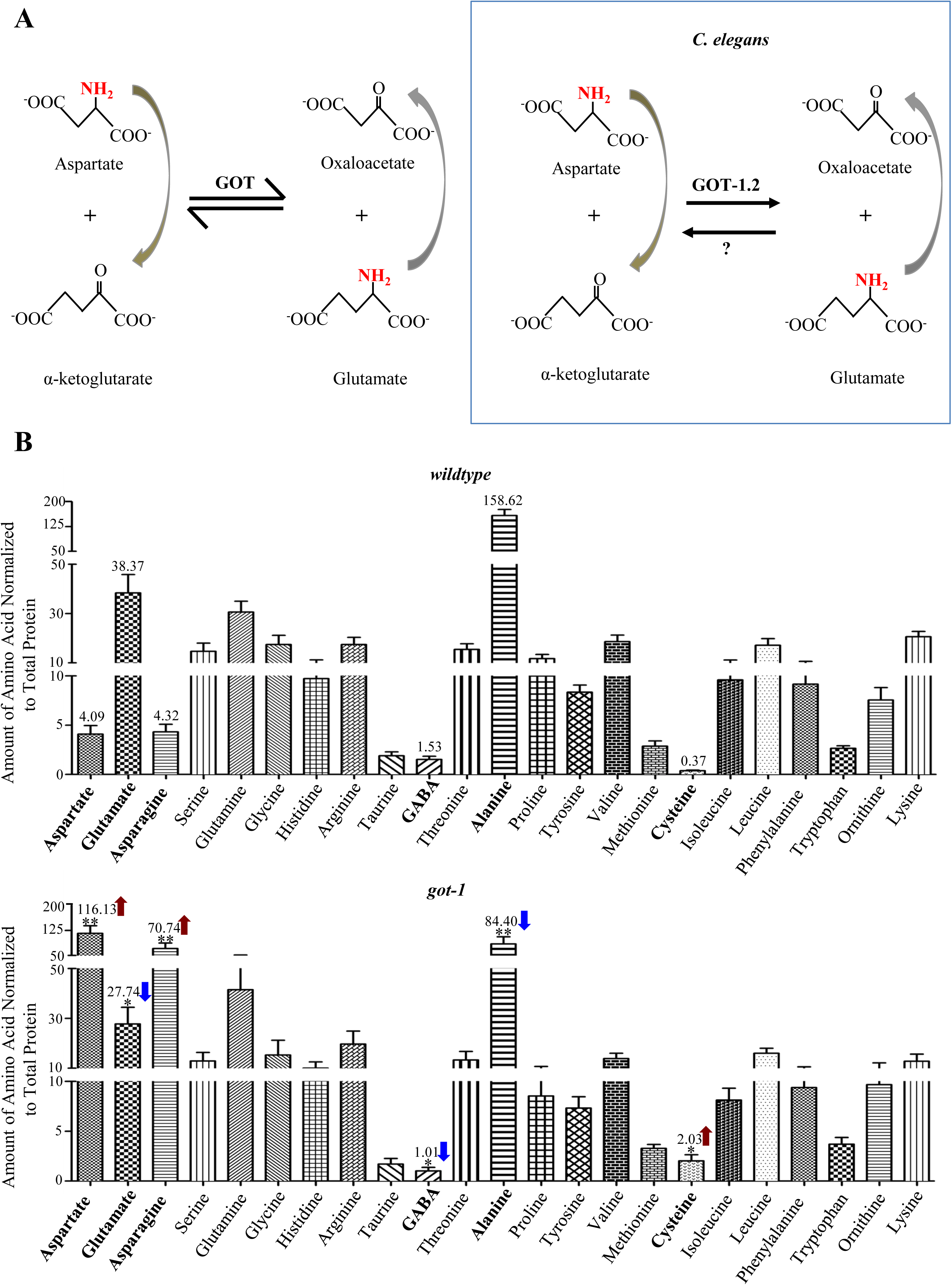
GOT-1 synthesizes glutamate from aspartate. A) GOT transaminases maintain the equilibrium between glutamate and aspartate by reversible transfer of the amino group (NH_2_) (left panel). Results from our amino acid profiling indicates that in *C. elegans*, GOT-1 preferentially catalyzes synthesis of glutamate from aspartate (right panel). B) Free amino acid levels measured from whole animal lysates by HPLC. *got-1* mutants (Bottom panel) exhibit reduced glutamate and alanine (blue arrows), and increased aspartate, asparagine and cysteine (red arrows) compared to wildtype animals (Top panel). In order to present data of a wide range, the Y-axis utilizes three different scales at different concentration values. Consistent with glutamate being the sole precursor of GABA synthesis, GABA level was also increased in *ubr-1* mutants (blue arrow). *P<0.05, **P<0.01 by the Student T test, N: 5 replica. Data are represented as mean ± SEM.

The genetic interaction between *ubr-1* and *got-1.2* is remarkably specific: *lf* mutations for other three GOT homologues did not restore bending of *ubr-1* mutants (Table 2). *lf* mutations in other metabolic enzymes or transporters that may be involved in glutamate and aspartate metabolic pathways (Table 2), including the alanine aminotransferase, glutamine synthetase, and glutaminase, did not rescue *ubr-1* mutant’s motor defects either. Henceforth, we refer to *got-1.2* as *got-1*.

**Table 2.** A list of genetic mutants in metabolic and other biological pathways examined for their genetic interactions with *ubr-1* mutants.

| Biological Pathways |  | Genes | Alleles (lf) | Background | Reversals |
| --- | --- | --- | --- | --- | --- |
| Glutamate Metabolism | Glutamate oxaloacetate transaminases | *got-1.2* | *hp731* | *ubr-1*<br>(*hp684*) | +++ |
|  |  |  | *hp731/mnDfl* |  | +++ |
|  |  | *got-1.1* | *tm2311* |  | - |
|  |  | *got-1.3* | *tm3424* |  | - |
|  |  | *got-2.1* | *gk109644* |  | - |
|  | glutamic acid decarboxylase | *unc-25* | *e156* |  | - |
|  | Glutamine synthetase | *gln-1* | *gk329791* |  | - |
|  |  | *gln-5* | *gk875298* |  | - |
|  | Glutaminase | *glna-2* | *gk536170* |  | - |
|  | alanine aminotransferase | *C32F10.8* | *tm2997* |  | - |
|  | GABA transaminase | *gta-1* | *ok517* |  | - |
|  | GABA transporter | *snf-11* | *ok156* |  | - |
| TCA Cycle | isocitrate dehydrogenase | *idhb-1* | *ok2368* |  | - |
|  | pyruvate dehydrogenase | *pdhk-2* | *tm3233* |  | - |
|  | pyruvate carboxylase | *pyc-1* | *tm3788* |  | - |
|  | pyruvate kinase | *pyk-1* | *ok1754* |  | - |
| Urea Cycle | carbamoyl phosphate synthetase 1 | *T28F3.5* | *tm4283* |  | - |
| mTORC | S6K | *rsks-1* | *ok1255* |  | - |
| Hetero-chrony | RNA binding | *lin-28* | *n719* |  | - |
|  |  |  | *n947* |  |  |
| Apoptosis | Caspase | *ced-3* | *n717* |  | - |
|  | CED-4 | *ced-4* | *n1162* |  | - |

### A critical requirement of GOT-1 activity in premotor interneurons to affect bending

To determine the endogenous expression pattern of GOT-1, we generated a functional GOT-1 reporter by inserting GFP at the endogenous *got-1* locus. GOT-1::GFP exhibited cytoplasmic expression in all somatic tissues, including broad expression in the nervous system (Fig. 4B; Fig. S2B). To determine the critical cells that mediate the genetic interactions between *ubr-1* and *got-1*, we restored cell-type specific GOT-1 expression in *ubr-1; got-1* mutants, and assessed their effect on the animal’s motor pattern. Restoring GOT-1 in all neurons (*Prgef-1*), but not in all muscles (*Pmyo-3*), fully reverted the motor pattern of *ubr-1; got-1* to the reduced bending as in *ubr-1* mutants (Table 1); therefore, both UBR-1 and GOT-1 function through neurons to regulate motor patterns.

Because GOT-1 is more broadly expressed in the nervous system than the UBR-1::GFP reporter (Fig. 4B), we examined whether GOT-1 functions through UBR-1-expressing neurons to regulate bending. Similar to our observation for neuronal sub-type UBR-1 rescue (Fig. 2), restoring GOT-1 in premotor interneurons of the reversal circuit (Table 1) was required for reversion of *ubr-1; got-1*’s bending pattern to that of *ubr-1*. Similarly, restoring GOT-1 in the same subset of these premotor interneurons, including AVE and RIM, exerted the most significant, partial reversion of *ubr-1; got-1*’s motor defects (Fig. 4D; Fig. S4).

Therefore, not only does a predicted metabolic enzyme GOT-1 exhibit genetic interaction with UBR-1, but also both proteins exhibit similar prominent requirement in premotor interneurons to regulate the reversal motor pattern. These results raise the possibility that metabolic dys-regulation may underlie *ubr-1*’s motor defects.

### GOT-1 synthesizes glutamate using aspartate

The catalytic activity of GOT-family transaminases enables reversible conversions between aspartate and glutamate (left panel in Fig. 5A). Their *in vivo* activity and physiological function, however, have not been examined in animal models.

To determine the metabolic changes associated with the loss of GOT-1, we uantified the amino acid levels of synchronized wildtype and *got-1* adults by highperformance iquid chromatography (HPLC). As previously reported (50), glutamate is an abundant amino acid, whereas aspartate is maintained at a low abundance in *C.elegans* upper panel in Fig. 5B). In *got-1* mutants, glutamate level exhibited a decrease of 27.6 ± 7.0% (mean ±SEM, n=4, P=0.0477 against wildtype animals), whereas the aspartate level exhibited a massive increase of 27.3 ± 4.76 folds (n=4, P=0.0076 against wildtype animals) (lower panel in Fig. 5B; Fig. 6A, B). These results implicate that *in vivo*, GOT-1 preferentially synthesizes glutamate from aspartate (right panel in Fig. 5A). Among the *C. elegans* GOTs, GOT-1 appears to be the key glutamate-synthesizing enzyme, because removing its homologue, GOT-2, did not lead to glutamate reduction (Fig. 6A).

**Figure 6.**
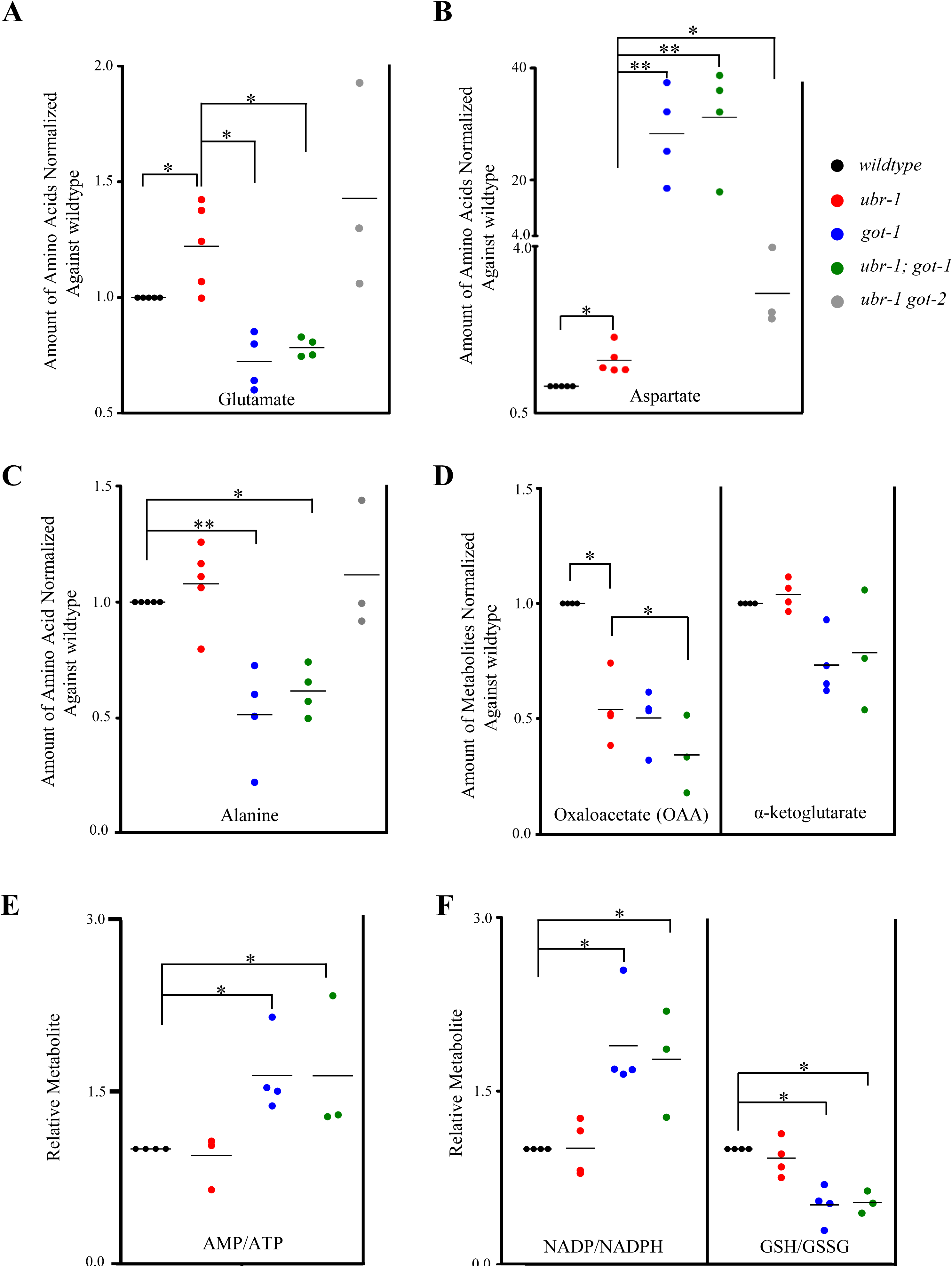
Glutamate level is elevated in *ubr-1* mutants. A-C) Free amino acid measured by HPLC, normalized against total protein in the lysate. Levels in mutants were normalized to that of wildtype. A) Glutamate level was increased in *ubr-1*, but decreased in *got-1* mutants; the increase in glutamate was reversed in *ubr-1; got-1*, but not in *ubr-1 got-2* mutants. B) Aspartate exhibited a modest increase in *ubr-1*, but massive accumulation in *got-1* and *ubr-1; got-1* mutants. The Y-axis utilizes two different scales (0.5 to 4.0, 4.0 to 40, respectively) to accommodate vastly different values. C) Alanine was significantly decreased in both *got-1* and *ubr-1; got-1* mutants. D-F) Metabolites measured by LC-MS/MS. D) OAA were significantly decreased in all mutants; α-KG did not exhibit consistent changes. E) The ratio of AMP to ATP was increased in *got-1* and *ubr-1; got-1* whereas *ubr-1* mutants did not show significant changes. F) The ratio of NADP to NADPH was increased in *got-1* and *ubr-1; got-1* mutants (left panel), while that of glutathione (GSH) to glutathione disulfide (GSSH) was decreased in both mutants (right panel). *ubr-1* mutants show no significant change in either ratio. *P<0.05, **P<0.01 by the Student T test. Horizontal lines represent mean values.

We noted that compared to the massive aspartate accumulation, the reduction of glutamate was mild in *got-1* mutants. This may result from compensatory activation of glutamate synthesis using other amino acids. As reported (50), alanine is the most abundant amino acid in *C. elegans* (upper panel in Fig. 5B). In *got-1* mutants, alanine level exhibited a decrease of 48.5 ± 12.4% (n=4, P=0.0050 against wildtype animals) (lower panel in Fig. 5B; Fig. 6C), supporting the notion that alanine becomes the compensatory source for glutamate when conversion from aspartate is blocked.

Such a drastic shift in equilibrium of the three key amino acids - aspartate, glutamate, and alanine - in *got-1* mutants must exert indirect metabolic consequences. To determine whether GOT-1’s loss affects the global metabolic state, we performed liquid chromatography-mass spectrometry (LC-MS) analyses on whole worm lysates. Indeed, *got-1* mutants exhibited increased AMP/ATP and NADP/NADPH, and decreased GSH/GSSG glutathione ratios (Fig. 6E, F), two hallmark features for increased cellular toxicity and metabolic stress (51).

We conclude that in *C. elegans*, GOT-1 synthesizes glutamate, and maintains glutamate level using aspartate. The loss of GOT-1 leads to glutamate reduction, aspartate accumulation, and potential compensatory glutamate synthesis using other amino acids.

### Glutamate level is elevated in *ubr-1* mutants

Removing the glutamate-synthesizing GOT-1 restored *ubr-1*’s bending pattern, suggesting that *ubr-1*’s motor defects may be associated with glutamate homeostasis.

We assessed the amino acid level in *ubr-1* mutants by HPLC. In *ubr-1* mutants, the glutamate level was increased by 22.2 ± 9.3% (n=5, P=0.0479 against wildtype animals). In *ubr-1; got-1* mutants, similar to *got-1* mutants, the glutamate level was reduced by 21.6 ± 2.4% (n=4, P=0.0230 against wildtype animals; P=0.0153 against *ubr-1* mutants), whereas the aspartate level was increased by 30.2 ± 5.3 folds (n=4, P=0.0033 against wildtype animals, P=0.0031 against *ubr-1* mutants) (Fig. 6A). Interestingly, there was a mild increase of aspartate in *ubr-1* mutants (46.3 ± 12.4%, n=4, P=0.0302 against wildtype animals) (Fig. 6B), which may reflect a compensatory metabolic response to reduce glutamate accumulation. Reminiscent of the genetic interactions exhibited at the behavioral level, removing GOT-1’s homologue, GOT-2, which did not restore *ubr-1*’s bending, did not reduce *ubr-1*’s glutamate either (Fig. 6A).

To determine if homeostasis of other metabolic substrates for GOT-1, α-KG and OAA, also contributes to *ubr-1*’s motor defects, we surveyed their levels in *ubr-1, got-1* and *ubr-1; got-1* adults by LC-MS/MS. Unlike the case for glutamate, which exhibited inversely correlated changes between *ubr-1* and *got-1* mutants, and between *ubr-1* and *ubr-1; got-1* mutants, the α-KG and OAA levels did not exhibit consistent, or correlated changes (Fig. 6C). These results support that restored bending in *ubr-1; got-1* animals primarily results from reduced glutamate, not from indirect - consequential or compensatory - metabolic effects that the reduced glutamate may have incurred.

### Some premotor interneurons exhibit defective morphology in *ubr-1* adults

Glutamate is a neurotransmitter, as well as an abundant amino acid partaking in other metabolic pathways including amino acid synthesis, energy production and urea cycle. An elevation of glutamate level may result in not only increased neuronal signaling in glutamatergic neurons, but also cellular stress in all UBR-1-expressing cells.

Because most of the critically required premotor interneurons for UBR-1’s role in motor pattern are cholinergic (52), the developmental effect of increased glutamate in these neurons may play a prominent role. To explore this possibility, we examined these interneurons using a transcriptional reporter (*Pnmr-1-*RFP) (18) that allows visualization of their somata (Fig. 7; Fig. S6), and a translation reporter (GLR-1::GFP) (18) that labels synapses of the AVA and AVE premotor interneurons(18) (Fig. 7).

**Figure 7.**
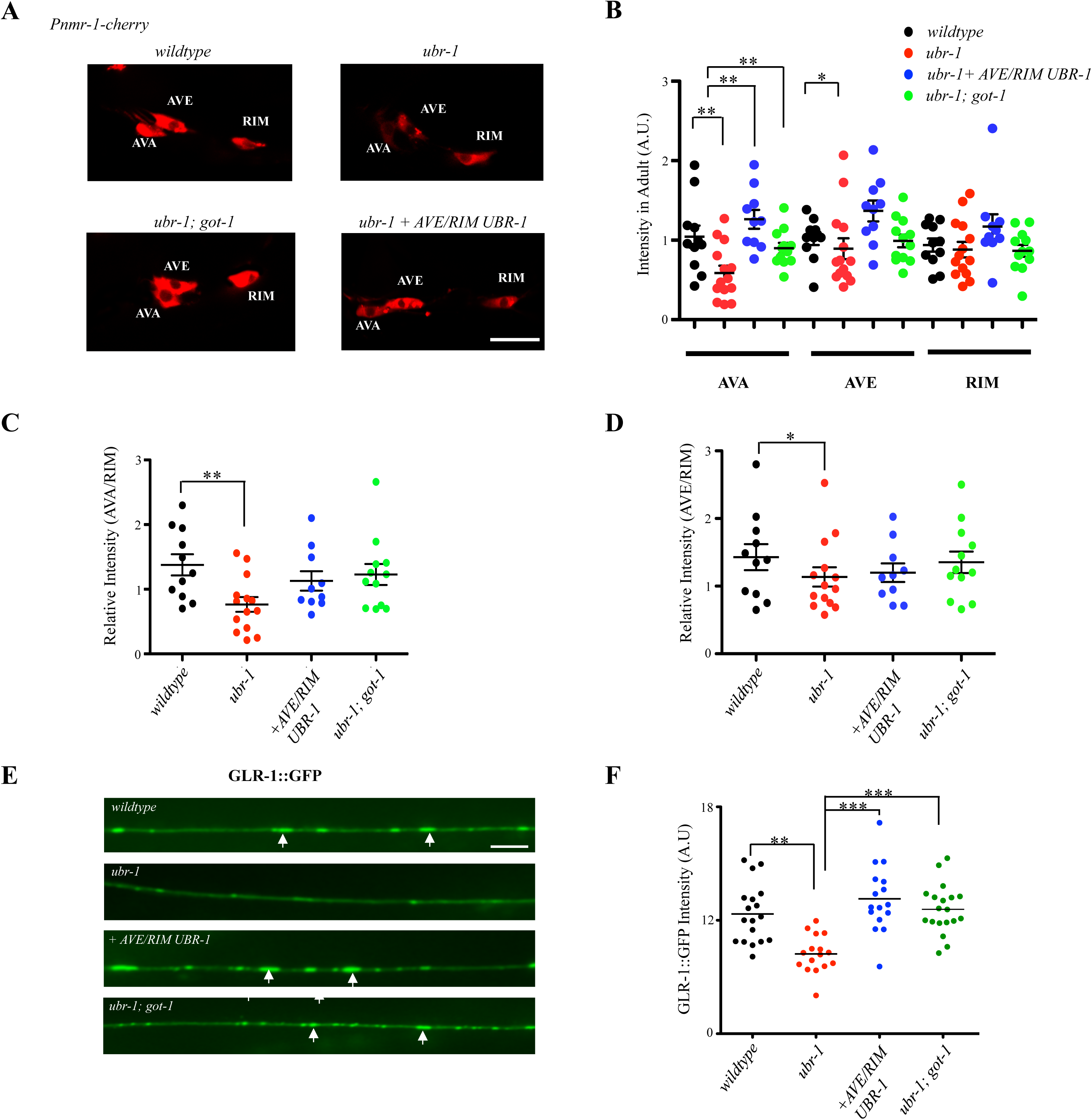
The loss of UBR-1 affects glutamate receptor expression in premotor interneurons AVA and AVE. A) Representative confocal images of AVA, AVE, and RIM neurons visualized with the *Pnmr-1-RFP* reporter in wildtype*, ubr-1,* and *ubr-1; got-1* adult animals, as well as in *ubr-1* mutants with restored expression of UBR-1 in neurons including AVE and RIM. Note the prominent reduction of fluorescent intensity in AVA in *ubr-1* mutants. B-D) Quantification of the fluorescent intensity in AVA, AVE and RIM (B), and relative fluorescent intensity between AVA and RIM (C) and between AVE and RIM (D), in animals with the respective genotypes denoted in A. N: more than 10 animals of each strain. * P<0.05, ** P< 0.01by the two-way ANOVA test. E) Representative confocal images of the GLR-1::GFP signals along the AVA and AVE neurites in the ventral nerve cords of animals with the same genotypes as denoted in panel A. *ubr-1* mutants exhibited a marked decrease in fluorescent intensity. F) Quantification of the total GLR-1::GFP intensity along the AVA and AVE ventral cord neurites. N=10-15, *P<0.05, **P<0.01,***P<0.001 by the Kruskal-Wallis test. Data are represented as mean ± SEM. Scale bar, 5 μm.

In adult *ubr-1* mutants, both reporters exhibited prominent and moderate decrease in fluorescence intensity in premotor interneurons AVA and AVE, respectively (Fig. 7A-D for P*nmr-1*; E-F for GLR-1::GFP). Such a decrease was not observed for other interneurons including RIM (Fig. 7A-D). We further noted that unlike the smooth and round somata in wildtype animals (Fig. S6, wildtype), in *ubr-1* mutants, the AVA and AVE somata exhibited rough edges and short branches in L4 stage larvae and adults (Fig. S6A, L4 and Adult panels), whereas other premotor interneurons such as RIM were appeared as in wildtype animals (Fig. S6B). The late onset of reversal bending defects coincides with the morphological change in AVA and AVE becoming prominent by the end of the larval stage (Fig. S6A). Their fluorescent intensity and morphology defects were significantly rescued by restoring UBR-1 expression in premotor interneurons (Fig. 7A-D; Fig. S6C), and in adult *ubr-1; got-1* mutants (Fig. 7A-D; Fig. S6C).

These results support the notion that an increased glutamate level may lead to developmental defects in *ubr-1* mutants, and premotor interneurons, in particular AVA and AVE, may be more susceptible to such a perturbation than other neurons or cells.

### GOT activity but not total protein level is increased in *ubr-1* mutants

Our analyses attributed the motor defect of *ubr-1* mutants to altered glutamate metabolism, most critically from premotor interneurons. To address how UBR-1 may negatively regulates glutamate level through GOT-1, we first assayed the total GOT activity of the whole worm lysates. We observed a moderate increase of GOT activity in *ubr-1* mutants (Fig. 8A). Consistent with the presence of multiple GOT homologues, the total GOT activity was drastically reduced, but not abolished in *got-1* mutants. The increased GOT activity in *ubr-1* mutants was attenuated in *ubr-1; got-1* double mutants. The attenuation effect was also specific to GOT-1: the loss of GOT-2 did not reduce GOT activity in *ubr-1* mutants (Fig. 8A).

**Figure 8.**
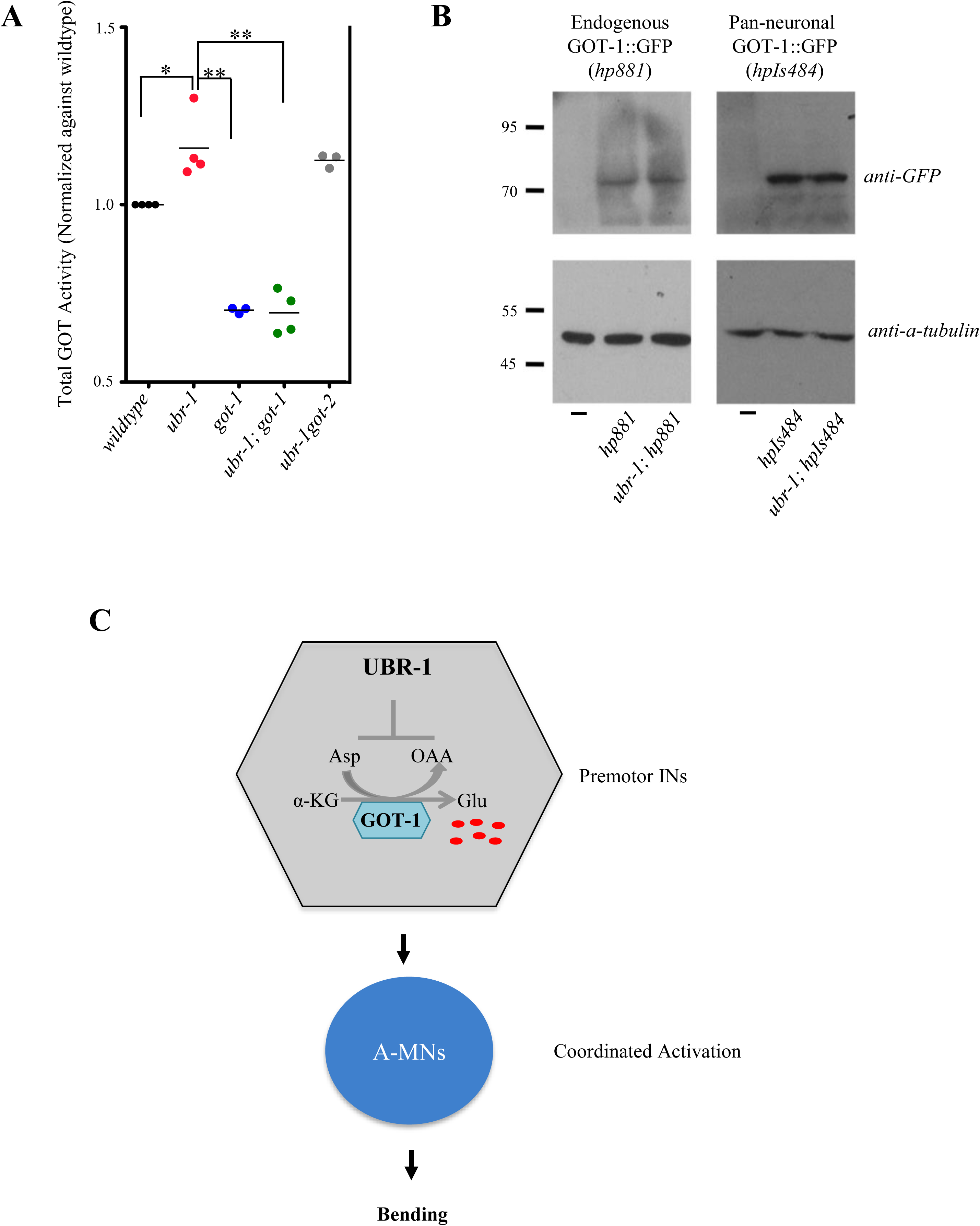
GOT-1 is not a direct substrate of UBR-1. A) The total GOT transaminase activity was increased in *ubr-1* and *ubr-1got-2* mutants, but decreased in *got-1* and *ubr-1; got-1* mutants. *P<0.05; **P<0.01 by the Kruskal-Wallis test. Horizontal lines represent the mean values. B) The loss of *ubr-1* did not lead to changes in GOT-1::GFP protein levels in *C. elegans* carrying endogenously fused GOT-1::GFP (left panel) or GOT-1::GFP driven by an exogenous pan-neuronal promoter (right panel). C) A working model of UBR-1-mediated negative regulation of glutamate level and bending.

An increased GOT activity is consistent with an elevated glutamate level in *ubr-1* mutants. However, we did not observe changes in GOT-1 protein level in *ubr-1* mutants. The level of GOT-1::GFP, expressed either from its endogenous locus or from an exogenous panneuronal promoter (*Prgef-1*), was similar between wildtype and *ubr-1* mutants (Fig. 8B). These results suggest that GOT-1 is not a direct substrate of UBR-1. UBR-1-mediated regulation of glutamate metabolism may involve targeting other pathway components.

## Discussion

Dys-regulated E3 activities have been implicated in neurodevelopmental and neurological disorders (53). Using the *C. elegans* model and its motor output as the functional readout, we reveal a previously unknown role of UBR-1 in glutamate metabolism. The absence of UBR-1 elevates glutamate level, which promotes simultaneous activation of A-class motor neurons, leading to reduced bending during reversal movements. This defect is compensated by removing a glutamate-synthesizing enzyme and reducing glutamate level in premotor interneurons of the reversal motor circuit. Aberrant glutamate metabolism may underlie *ubr-1* mutant’s physiological defects.

### *ubr-1*’s motor phenotype is associated with glutamate level change

Coinciding with an elevated GOT activity, *ubr-1* mutants exhibit increased glutamate level. Removing a key glutamate-synthesizing enzyme GOT-1 reduces the glutamate level in *ubr-1* mutants. These metabolic changes parallel those in motor behaviors: removing the activity of GOT-1 enzyme restores bending in *ubr-1* mutants. These results imply that elevated glutamate level is associated with *ubr-1*’s defective bending.

Genetic mutations in key metabolic enzymes can exert effects beyond their primary substrates and immediate metabolic pathways. For example, upon the loss of GOT-1, the reduction of glutamate synthesis from aspartate likely induces compensatory utility of other amino acids, thus the reduction of alanine. When converting aspartate to glutamate, GOT-1 may indirectly affect the equilibrium of other substrates, OAA and α-KG, metabolites that take part in multiple metabolic functions, including energy production, amino acid homeostasis, nucleotide synthesis, and lipid synthesis.

Our metabolomics analyses of *got-1* mutants revealed no consistent or correlated changes in the level of OAA, α-KG (Fig. 6C), and all other TCA cycle metabolites from wildtype controls (Fig. S5). Because the same samples exhibited consistent and correlated changes between the aspartate, glutamate and alanine levels, the variability in other metabolite levels likely reflects flexibility of compensatory or adaptive changes in response to the primary metabolic dysfunction upon the loss of GOT-1. Indeed, the only consistent and anti-correlated metabolic change that we observed between *ubr-1* and *got-1* (and *ubr-1; got-1*) mutants was the glutamate level. Cumulatively, these results strongly support that the primary metabolic change - the level of glutamate - causes the motor pattern change in *ubr-1* and *ubr-1; got-1* mutants.

Our analyses measured glutamate level in whole animals, not the specific neuron groups that are critically required for both UBR-1 and GOT-1’s effect on reversals. Examples of genes with broad expression, but cell-type specific functions have been reported (32, 54, 55). These and other results – such as the presence and requirement of both UBR-1 and GOT-1 in premotor interneurons in mediating their effects on the reversal motor pattern – suggest that the glutamate level change in premotor interneurons significantly contribute to the motor pattern change in *ubr-1* and *ubr-1; got-1* mutants. We reckon that other UBR-1-expressing neurons likely also contribute to such an alteration. Furthermore, the glutamate change in other neurons and cells may cause other physiological effects that were not examined in this study, where we only followed the most obvious motor defect.

### Increased glutamate may cause defective premotor interneuron development

Glutamate has multiple functions. Being a key neurotransmitter itself, glutamate also serves as the sole precursor of another neurotransmitter GABA. Glutamate is also one of the abundant amino acids partaking in other metabolic pathways including amino acid synthesis, energy production and urea cycle. An elevation of glutamate not only affects signaling in glutamatergic and likely GABAergic neurons, but also increases metabolic stress in all cells.

Because both UBR-1 and GOT-1 exert strong effects on the reversal motor pattern through premotor interneurons, most of which cholinergic (52), the developmental effect of increased glutamate may play a prominent role in these interneurons. Indeed, we observed morphological changes of premotor interneurons that coincided with the onset of motor pattern change in *ubr-1* mutants. Interestingly, within the reversal motor circuit, premotor interneurons that exhibited the prominent changes, AVA and AVE, provide direct synaptic inputs to the A-class motor neurons. These observations suggest that compared to other cells and neurons, the development and function of premotor interneurons, in particular AVA and AVE, may be more susceptible to glutamate increase, likely through both increased glutamatergic inputs and cellular stress.

### GOT-1 is a key enzyme for glutamate homeostasis in neurons

Despite its broad expression, GOT-1 can only influence *ubr-1*’s motor defects through neurons. These results indicate that maintaining glutamate level in neurons, especially the premotor interneurons, is critical for the proper motor output of *ubr-1* mutants.

In the mammalian brains, glutamate has to be locally synthesized because it cannot cross the blood-brain barrier (56). The activity of transaminases, mainly by GOT and ALT, establishes and maintains homeostasis of three abundant amino acids, glutamate, alanine, and aspartate (57, 58). A recent study suggests that GOT also contributes to glutamate synthesis at synapses (59).

The main sources of glutamate synthesis and homeostasis in *C. elegans* neurons have remained elusive (60). Our study establishes GOT-1 as a key enzyme for glutamate synthesis. Further, it provides the first evidence of a critical role of GOT-1 in glutamate metabolism in the nervous system.

### Direct substrates through which UBR-1 regulates glutamate level are unknown

How UBR-1 negatively regulates glutamate level remains to be elucidated. UBR family proteins are often addressed in the context of N-end rule E3 ligases (6, 7, 10). However, UBR proteins have non-N-end rule substrates, and N-end rule substrates do not account for all described functions of UBR proteins (21, 61-64), and their physiological relevance remains to be clarified.

For example, in mammalian cells, UBR1’s N-end rule substrates include pro-apoptotic fragments (11). In *C. elegans,* the CED-3 caspase promotes apoptosis. During postembryonic development, it interacts with UBR-1 to expose the N-degron of LIN-28 (10), a regulator of seam cell patterning (65) to promote its degradation. However, neither UBR-1 nor LIN-28’s N-degron were essential for LIN-28’s degradation (10). Removing either apoptotic regulators CED-3 or CED-4, which generates pro-apoptotic fragments, or LIN-28, neither mimic nor rescue *ubr-1* mutants’ bending defects (Table 2; Supplementary Movie 3). Substrates for neuronal function of UBR-1 remain to be identified.

Regardless of the nature of UBR-1 targets, our results show that UBR-1 is unlikely to directly regulate GOT-1’s protein stability: we did not observe a change in the level of GOT-1::GFP in *ubr-1* mutants. We speculate that UBR-1-mediated regulation of glutamate level may involve targeting other components that subsequently affect glutamate metabolism. For example, the activity of many rate-limiting metabolic enzymes in glycolysis, fatty acid synthesis, cholesterol synthesis, and gluconeogenesis are regulated by phosphorylation and de-phosphorylation (66, 67). UBR-1 may mediate the ubiquitination of kinases or phosphatases that modify GOT-1’s activity. A large-scale *ubr-1* suppressor screen may be required to yield insights on UBR-1’s direct substrates.

### The *C. elegans* motor behavior as a model to investigate UBR1 and *JBS*

The UBR family proteins have been extensively examined in yeast, cultured cells, and mouse models, but only recently in *C. elegans*. In yeast, UBR1 affects cell cycle progression, but is non-essential (6). Mouse models revealed the functional redundancy of multiple UBR homologues, where combinatorial knockout of UBR1 and UBR2 results in embryonic lethality (12). *C. elegans* UBR1 affects the stability of LIN-28, a regulator of postembryonic hypoderm seam cell division (10); both UBR1 and LIN-28 are non-essential for viability. The simplicity and viability of the *C. elegans ubr-1* model has allowed us to reveal, and genetically dissect a previously unknown physiological role of UBR-1 in glutamate homeostasis. We note that aberrant glutamate metabolism may cause systemic cellular and other unexamined neuronal defects in *ubr-1* mutants. However, the simplicity and sensitivity of the *C. elegans* premotor interneuron circuit, and the prominent motor pattern change provided a quantifiable functional readout to afford genetic pathway dissection.

Our study provides the first demonstration of a UBR protein’s physiological role in glutamate regulation. We noted with interest that in a case study of a *JBS* patient with severe cognitive impairments, GOT activity levels, used as a biomarker of inflammation, were increased (68). In addition to its role in development, there is a growing body of evidence for glutamate signaling in non-neuronal tissues (48, 69, 70). It will be of interest to examine whether glutamate level and signaling in other *JBS* animal models and *JBS* patients are aberrant, and whether they contribute to *JBS* pathophysiology.

## Methods

### Strains and constructs

#### Strains

See Tables 1, 2 and S1 for a complete strain list generated in this study. All *C.elegans* were cultured on standard NGM plates seeded with OP50, and maintained at 22°C.

*hp684* and *hp731* were isolated by EMS mutagenesis. They were mapped using SNP mapping and whole genome sequencing (71, 72), followed by behavioral rescue using fosmids and genomic fragments that harbor *ubr-1* and *got-1.2*, respectively. Placing *hp731* over a deficiency *mnDf1* fully recapitulated the rescuing effect in the *ubr-1* mutant background, confirming that *hp731* being a *lf* allele of *got-1.2* (Table 2)*. hp820, hp821, hp820hp833* and *hp865* were generated by CRISPR-Cas9-mediated genome editing, following protocols described in (26). The rest of genetic mutants were obtained from *CGC*; all strains were backcrossed at least four times against N2 by genotyping.

Transgenic strains include those with extra-chromosomal multi-copy arrays (*hpEx*), integrated multi-copy arrays (*hpIs)*, single-copy integrated arrays (*hpSi*), and a *got-1.2* GFP knock-in allele (*hp*). Transgenic animals carrying extra-chromosomal arrays (*hpEx*) were created by co-injecting the DNA plasmid of interest and a co-injection marker at 5-30 ng/μL. Extra-chromosomal arrays were integrated into the genome using UV irradiation to create stable, transgenic lines (*hpIs*) (73). We found that GOT-1 overexpression rendered sick animals. Hence, all *got-1* tissue-specific rescue lines were generated using Mos1-mediated single copy insertion (*hpSi*) as previously described (74). All integrated strains were outcrossed several times against N2 prior to phenotypic analyses.

#### GOT-1.2::GFP knock-in

The in-frame GOT-1 C-terminal GFP fusion allele (*hp881*) was generated by CRISPR-Cas9 mediated homologous recombination, following protocols described in (75). The replacement template for GOT-1::GFP (pJH3629) includes 1kb sequence upstream to the *got-1* start codon, the entire *got-1* coding sequence, and 1.5kb sequence downstream of the *got-1* stop codon. GFP was inserted in-frame after the last amino acid of GOT-1. A LoxP-Prps-27-NeoR-loxP cassette was inserted between the stop codon and 3’ UTR. After injection, animals were allowed to lay eggs overnight at 25^^0^^C before G418 selection. Animals with extra-chromosomal arrays were selected against as described. Candidates for insertion were confirmed by genotyping and sequencing. Confirmed insertion lines were injected with a *Pelt-3::*Cre plasmid to remove the LoxP cassette. The resulting *hp881* allele was outcrossed against N2 wildtype animals twice before crossed into the *ubr-1* mutants.

### Locomotion Analysis

#### Plate conditions

When transferred to a new, thinly seeded plate, *C. elegans* typically spend most of the time moving forward, with brief interruptions of backward movement. As previously described (42), 35 mm NGM plates with limited food (lightly seeded OP50 bacteria) were used for automated tracking and behavioral analyses. Here we quantified body curvature, initiation frequency, and duration on one-day old young adults using copper chloride (CuCl_2_), an aversive stimulus for *C. elegans* (76). Briefly, immediately before transferring worms onto the NGM plates, a ∼10-20 μl of 100 mM CuCl_2_ solution was pipetted into a small circle (∼20 mm in diameter) in the middle of the plate for the animal to roam. One animal was placed in the center of the circle and allowed to habituate for one minute prior to recording for three minutes. For the next recording,CuCl2 was pipetted onto the same circle before another animal was placed on the plate and recorded. For all data presented in the same graph, animals were recorded on the same day, with at least one animal from each genotype recorded on the same plate. We altered the order of recording for animals of different genotype.

#### Tracking and data analysis

Behavior was recorded using a Zeiss Axioskop 2 Plus equipped with an ASI MS-40000 motorized stage and a CCD camera (Hamamatsu Orca-R2). Tracking and analysis were performed using Micromanager and Image J software plug-ins developed in-house (courtesy of Dr. Taizo Kawano). Image sequences were sampled at 100-msec exposure (10 frames per second). For the post-imaging analysis, an Image J plug-in was used to skeletonize the worm and extract its centerline. The centerline was divided into 29 equal segments, and the angle between each segment and its tangent line was calculated to quantify the curvature of the animal. These segments were binned into four groups to capture the most consistent curvature trends across the entire length of the animal. The directionality of movement (forward vs. backward) was determined by first identifying the anterior-posterior axis or the “head” and “tail” points, which was manually defined at the first two frames and verified throughout the recording. To calculate directionality of movement, the displacement of the midline point in relation to the head and tail for each worm was determined based on its position in the field-of-view and the stage coordinates. Image sequences wherein animals touched the edge of the recording field and crossed over on themselves were not processed.

#### Quantification

Analyses of the output data were carried out using an R program-based code developed in-house (courtesy of Dr. Michelle Po). From a group of recordings (N≥10 for each genotype per experiment), we quantified the following parameters,among other behavioral parameters analyzed by the program: 1) Curvature, the average curvature of each animal at each segment; 2) Initiation (defined as the frequency of directional change for each animal; 3) Duration (defined as the time spent moving in the same direction for >3 frames or 300msec, calculated for each bout of forward or backward initiation). Body curvature, initiations, and durations were calculated for forward and backward locomotion separately.

### The A-class motor neuron calcium imaging and analysis

Animals of various genotypes were crossed into a reporter *hpIs460* (*Punc-4-GCaMP6::wCherry*), using a calcium reporter GCaMP6 fused in-frame at its C-terminus with wCherry for both tracking and ratiometric correction for motion artifacts in moving animals during recordings (46). A-MN imaging in moving young adults was carried out as described previously (32). Briefly, *C. elegans* were placed on a 2.5% agar pad, immersed in a few drops of M9 buffer, and covered by a coverslip to allow slow crawling. Each recording lasted for 3 minutes. Images were captured using a 40x objective on a Zeiss Axioskop 2 Plus equipped with an ASI MS-40000 motorized stage, a dual-view beam splitter (Photometrics) and a CCD camera (Hamamatsu Orca-R2). The fluorescence excitation light source from X-CITE (EXFO Photonic Solution Inc.) was reduced to prevent saturation of imaging field. The fluorescent images were split by Dual-View with a GFP/RFP filter set onto the CCD camera operated by Micromanager. The 4x-binned images were obtained at 50-msec exposure time (10 frames per second). Regions of interest (ROIs), containing the soma of DA7, VA10 and VA11, were defined and simultaneously tracked using an in-house developed Image J plug-in (32). The ratio between GFP and RFP fluorescence intensities from individual ROI was plotted over time to produce index for calcium profile for each neuron. Displacement for the DA7 soma, which exhibited the strongest RFP signal, was plotted over time to generate the example instantaneous velocity profiles. For example velocity profiles, we also manually annotated videos to verify the reversal periods. Reversal events longer than 5 seconds were used for cross-correlation analyses shown in the example traces.

The pair-wise phase relationships between the activity patterns of A-MNs during each reversal event were assessed by cross-correlation analyses. In each example trace, cross-correlation during the entire period of reversal was calculated to determine the phase lags between the calcium signals of VA11, DA7, and VA10. The maximum of the cross-correlation function denotes the time point when the two signals are best aligned; the corresponding argument of the maximum correlation values denotes the lag between the two neurons. The absolute time lags were used to represent the synchrony of VA11, DA7 and VA10’s activation. N (≥10) refers to the number of reversal events analyzed for each genotype per experiment.

### *C. elegans* metabolomics analyses by HPLC and LC-MS/MS

Synchronized last-stage larval worms were grown on 100 mm NGM agar plates seeded with OP50 bacteria. Worms were collected using M9 buffer and were washed thoroughly to remove bacteria. Worm pellets were snap-frozen in liquid nitrogen and pulverized using a cell crusher. Amino acids and other metabolites were extracted by addition of ice-cold extraction solvent (40% acetonitrile, 40% methanol, and 20% water) and incubated on dry ice for an hour with occasional thawing and vortexing. Samples were then moved to a thermo mixer (Eppendorf) and shook for an hour at 4**°**C at 1400 rpm. These samples were centrifuged at 14000 rpm for 10 min at 4**°**C. Supernatant was transferred to fresh tubes and lyophilized in a CentreVap concentrator (Labconco) at 4**°**C. Samples were stored at **-**80 **°**C until used for HPLC or LC**-**MS/MS analyses.

Amino acid quantitation was performed using the Waters Pico-Tag System (Waters). After hydrolysis and pre-column derivatization of the sample by PITC, samples were analyzed by reverse phase HPLC (Amino acid facility, SPARC BioCentre, Sick Kids, Toronto, Canada). LC-MS/MS metabolite analysis was performed as described previously (77). Raw values were normalized against the total protein concentration as determined by a Bradford Protein Assay (Bio-Rad). Results were compared from at least three sets of independent experiments (N≥3), with all samples collected and analyzed in parallel in each replica.

### Measurement of total GOT Activity from the *C. elegans* lysate

Synchronized late L4 larval stage worms were grown on 100 mm NGM agar plates seeded with OP50 bacteria. Worms were collected using M9 buffer and were washed thoroughly to remove bacterial contamination. Samples were homogenized by sonication in 100-200ul ice-cold AST buffer. Samples were centrifuged at 13,000*g* for 10 minutes to remove insoluble materials. The supernatant was used for the AST assay and pellets were used to quantify the total protein in the samples (as described above).

GOT (also referred to as AST) activity was measured using a colorimetric assay with AST Activity Assay Kit (Sigma-Aldrich) according to the manufacturer’s instructions. Data were normalized by the total protein content of the whole worm lysate as determined by a Bradford Protein Assay.

### *C. elegans* Biochemistry

Mixed stage *C. elegans* were grown on 100 mm NGM plates seeded with OP50 and collected using M9 buffer. Lysates were prepared as described previously (78). For western blot analyses, total protein concentration was determined using a Bradford Protein Assay (Bio-Rad). Anti-GFP antibodies (Roche) were used to probe for GOT-1::GFP and tubulin was used for the loading control.

### Statistical Analysis

For bending curvature and calcium imaging analyses, statistical significance was determined using the Kruskal-Wallis test and the two-way repeated measures (RM) ANOVA and subsequent post-hoc analysis. For metabolite analyses, two-tailed Student’s t-tests were applied to determine statistical differences. p< 0.05 were considered to be statically significant. All statistics were performed using Prism software (GraphPad).

## Acknowledgements

We thank Asuka Guan, Josh Kaplan, Gene Knockout Consortium and National Bioresource Project and *Caenorhabditis elegans Center* (CGC) (funded by NIH Office of Research Infrastructure Programs P40 OD010440) for strains, Reynaldo Interior for the HPLC analyses, Don Moerman and Stephen Flibotte for the whole-genome sequencing analyses, Amy Caudy for advice on the metabolomics analyses. We thank Taizo Kawano, Michelle Po, Valeriya Laskova, and Asuka Guan for technical and programming support. We thank Aravinthan Samuel and Andrew Chisholm for comments on the manuscript.

## Supporting Information

### Supplementary figure legends

**S1 Fig. *ubr-1* mutants exhibit reduced bending.**

A) Representative images of wildtype animals and two alleles of *ubr-1* mutants during reversals (left to right panels). The wildtype animal generates sinusoidal body bends, whereas *ubr-1(hp820, hp821)* animals do not bend. This defect was rescued by restoring the expression of UBR-1. Dots denote position of tail. Scale bar: 200µm. In *ubr-1(hp820)* (B) and *ubr-1(hp821)* (E) alleles (grey line), bending curvature is reduced throughout head to tail compared to wild type (black line), and this was rescued by restoring expression of UBR-1 using its endogenous promoter (red line). *ubr-1* mutants have fewer initiations (C, F) and longer durations for reversals (D, G). ***P<0.001, **P<0.01, *P<0.05 by the Two-way RM ANOVA test. Data are represented as mean ± SEM. ***P< 0.001, **P<0.01, *P< 0.05 by the Kruskal-Wallis test. Data are represented as mean ± SEM.

**S2 Fig. UBR-1 and GOT-1 are expressed in AVA, AVE and RIM**.

A) A confocal image of animals expressing a functional UBR-1::GFP transgene from its endogenous promoter (green), and the *Pnmr-1*-RFP reporter (red). B) A confocal image of animals expressing endogenous GOT-1::GFP from the knock-in allele (green), and the *Pnmr-1*-RFP reporter (red). Robust expression of UBR-1 and GOT-1 was present at the pharynx; neuronal UBR-1::GFP and GOT-1::GFP signals were present in the AVA, AVE and RIM premotor interneurons (denoted), and other unidentified neurons (not shown). Scale bar, 5 μm.

**S3 Fig. *ubr-1* function is required in premotor interneurons to regulate reversals.**

A-B) In *ubr-1* mutants, the reversal duration is increased, while reversal initiation frequency is decreased. Both the parameters were rescued by restoring the expression of UBR-1 in premotor interneurons, but not in GABAergic or cholinergic motor neurons. C-D) Expression of UBR-1 in multiple premotor interneurons, which include AVE/RIM exhibited significant rescue of *ubr-1* for reversal duration and initiation frequency. Expression of UBR-1 in the RIM alone or in AVA alone did not result in rescue.*P<0.05, **P<0.01, ***P<0.001 by the Kruskal-Wallis test. Data are represented as mean ± SEM.

**S4 Fig. GOT-1 function in premotor interneurons is required to suppress *ubr-1’s* reversal defects.**

The functional loss of *got-1* restores the reversal duration (A) and the reversal initiation frequency (B) in *ubr-1* mutants. Restoring GOT-1 expression in multiple premotor interneurons, including AVE and RIM, reverted the reversal duration of *ubr-1; got-1* to that of *ubr-1* (A), whereas the reversal initiation frequency exhibited the trend of decrease but did not reach statistic significance (B). Expression of GOT-1 in RIM alone had no effect, while the expression of GOT-1 in AVA partially reverted the reversal duration.*P<0.05, **P<0.01, ***P<0.001 by the Kruskal-Wallis test. Data are represented as mean ± SEM.

**S5 Fig. Metabolites of the TCA cycle in *ubr-1, got-1* and *ubr-1; got-1* mutants did not exhibit consistent changes from wildtype animals.**

Metabolite levels measured by LC-MS, normalized against the total protein level in the lysate. All mutants were normalized to that of wildtype animals. Metabolites of the TCA cycle did not show any coordinated changes among these mutants.

**S6 Fig. The AVA and AVE somata exhibit morphological changes in *ubr-1* mutants by the end of larval stage development**

A) Confocal images of the AVA and AVE premotor interneuron somata in wildtype and *ubr-1* animals, which were labeled with cytosolic RFP, from the L1 juvenile larva to adult stages. Depending on the focal planes, some images contain that of nuclei, which were devoid of RFP signals. Left panels: images from wildtype animals. The surface of both somata was round and smooth throughout larval development and in young adults. Right panels: images from *ubr-1* mutants. In young (L1 to L3) larva, somata appeared similar to those in age-matched wildtype animals. In the L4 larva and adult animals, somata developed rough surface and short branches, denoted by arrows. B) The RIM soma exhibited normal morphology in adults, similar to wildtype animals. C) The round morphology AVA and AVE somata in *ubr-1* adults were restored in *ubr-1; got-1* double mutant adults, and in *ubr-1* adults with restored UBR-1 expression in multiple premotor interneurons including AVE and RIM. Asterisks (*) mark the AVA or AVE axons, which could not be shown in some panels when they were in a different focal plane as the somata. Scale bar, 2μm (for the L1 and L2 panels) 5 μm (for the L3 to adult panels).

### Supplementary movie captions

**S1 Movie: Reversal behaviors exhibited by the wildtype (N2) (Part 1) and *UBR-1(hp684; lf)* (Part 2) young adults.**

The *ubr-1* mutant exhibits prominent rigidness during reversals.

**S2 Movie. Reversal behaviors exhibited by *ubr-1(hp684; lf)* (Part 1), *got-1(hp731; lf)* (Part 2), and *ubr-1(hp684; lf); got-1(hp731; lf)* (Part 3) young adults.**

The functional loss of GOT-1.2 in the *ubr-1* mutant background led to significant improvement of body bending during reversals.

**S3 Movie. Reversal behaviors exhibited by *lin-28(n719; lf)* (Part 1), and *UBR-1(hp684; lf) lin-28(n719; lf)* (Part 2) young adults.**

The functional loss of the LIN-28 protein in the *ubr-1(hp684; lf)* mutants did not improve body bending during reverses.

**Table S1.**
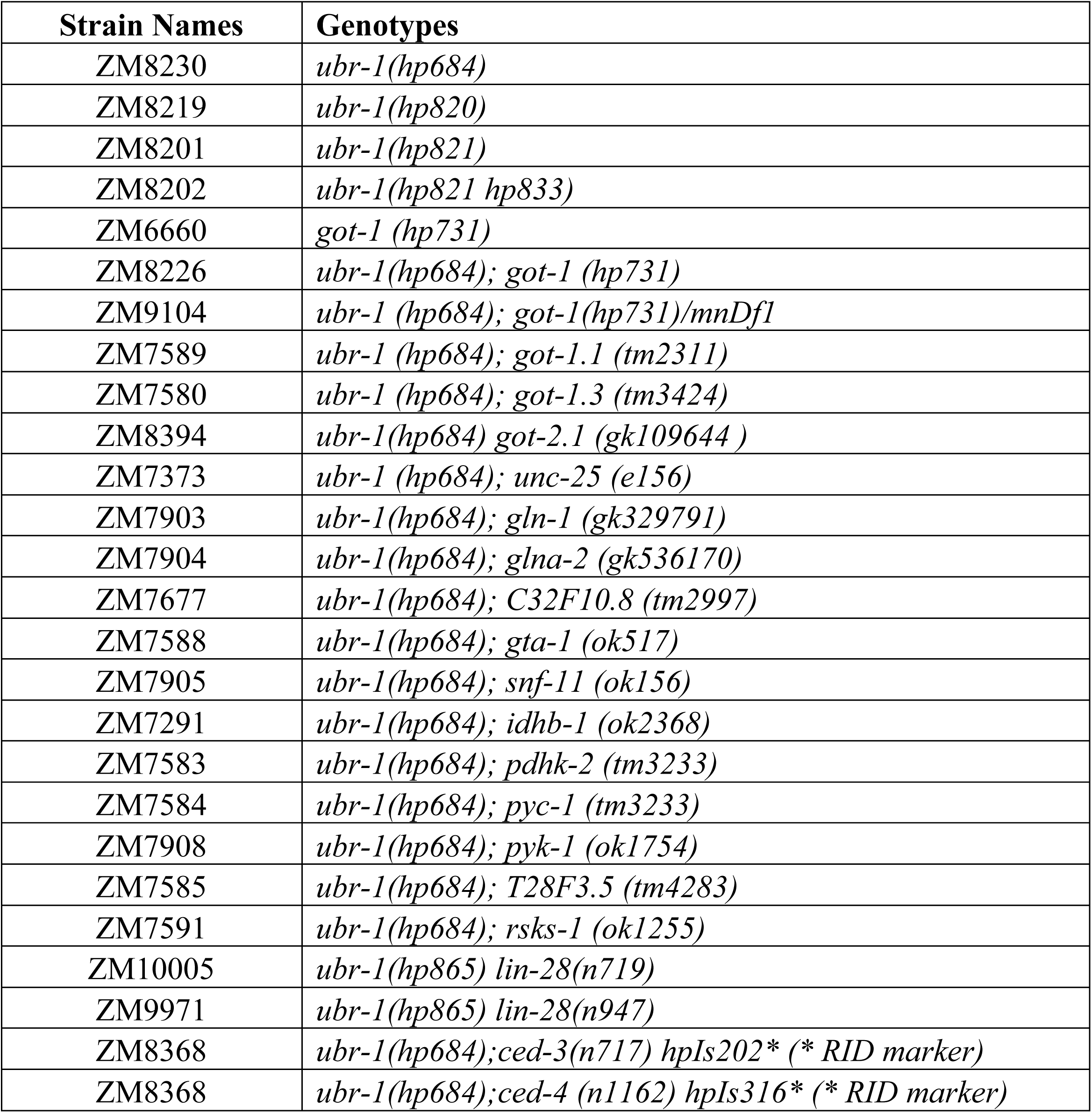
Strains generated or acquired for this study. **a. Non-transgenic strains generated for genetic interactions with *ubr-1***

**b. Transgenic strains generated for A-MN calcium imaging**

**b. Transgenic strains generated for A-MN calcium imaging**
| Strain Names | Genetic Backgrounds | Transgenes | Plasmid Names (Description) |
| --- | --- | --- | --- |
| ZM8426 | - | *hpIs460* | pJH3137 (*Punc-4-GCaMP6s::wCherry*) |
| ZM8749 | *ubr-1 (hp684)* |  |  |
| ZM8750 | *ubr-1 (hp684); got-1 (hp731)* |  |  |
| ZM8846 | *ubr-1(hp684)* | *hpIs460* | pJH3137 (*Punc-4-GCaMP6s::wCherry*) |
|  |  | *hpEx3676* | pJH3476 (*Popt-3::UBR-1 minigene*) |

**c. Transgenic strains generated for behavioral rescue, gene expression, and biochemistry experiments.**

1 c. Transgenic strains generated for behavioral rescue, gene expression, and biochemistry 2 experiments.
| Gene | Strain | Backgrounds | Trans-<br>genes | Plasmids |
| --- | --- | --- | --- | --- |
| U<br>B<br>R | ZM7440 | *ubr-1(hp684)* | *hpEx3161* | pJH2913 (*Pubr-1::UBR-1*) |
|  | ZM8272 | *ubr-1(hp821hp833)* | *hpEx3482* |  |
|  | ZM8270 | *ubr-1(hp820)* | *hpEx3480* |  |
|  | ZM8271 | *ubr-1(hp821)* | *hpEx3481* |  |
|  | ZM7410 | *lin-15(n765)* | *hpEx3056* | pJH2963 (*Pubr-1::GFP*) |
|  | ZM8628 | - | *hpIs487* | pJH2965 (*Pubr-1::GFP::UBR-1*) |
|  | ZM8389 | *ubr-1(hp684)* | *hpEx3535* | pJH3380 (*Prgef-1::UBR-1 minigene*) |
|  | ZM8388 |  | *hpEx3536* | pJH3381 (*Pmyo-3::UBR-1 minigene*) |
|  | ZM8390 |  | *hpEx3537* | pJH3382 (*Punc-47::UBR-1 minigene*) |
|  | ZM8391 |  | *hpEx3538* | pJH3389 (*Pacr-2::UBR-1 minigene*) |
|  | ZM8404 |  | *hpEx3541* | pJH3390 (*Pglr-1::UBR-1 minigene*) |
|  | ZM8457 |  | *hpEx3564* | pJH3436 (*Pnmr-1::UBR-1 minigene*) |
|  | ZM8551 |  | *hpEx3595* | pJH3469 (*Prig-3::UBR-1 minigene*) |
|  | ZM8550 |  | *hpEx3596* | pJH3476 (*Popt-3::UBR-1 minigene*) |
|  | ZM8582 |  | *hpEx3597* | pJH3485 (*Plgc-55s::UBR-1 minigene*) |
|  | ZM8627 |  | *hpEx3622* | pJH3502 (*Pgcy-13::UBR-1 minigene*) |
|  | ZM8580 |  | *hpEx3599* | pJH3487 (*Pttx-3::UBR-1 minigene*) |
|  | ZM8581 |  | *hpEx3598* | pJH3486 (*Pinx-1::UBR-1 minigene*) |
| G<br>O<br>T | ZM7289 | *TTti5605* | *hpSi10* | pJH2998 (*GOT-1.2::GFP MosCI*) |
|  | ZM8458 |  | *hpSi16* | pJH3396<br>(*Prgef-1::GOT-1.2 MosCI*) |
|  | ZM8460 |  | *hpSi18* | pJH3397<br>(*Pglr-1::GOT-1.2 MosCI*) |
|  | ZM8462 |  | *hpSi20* | pJH3399<br>(*Punc-47::GOT-1.2 MosCI*) |
|  | ZM8497 |  | *hpSi22* | pJH3471 (*Prig-3::GOT-1.2 MosCI*) |
|  | ZM8760 | *ubr-1(hp684);*<br>*got-1(hp731)* | *hpEx3661* | pJH3531 (*Popt-3::GOT-1.2*) |
|  | ZM8759 |  | *hpEx3621* | pJH3501 (*Pgcy-13::GOT-1.2*) |
|  | ZM8102 |  | *hpEx3998* | pJH2896 (*Pmyo-3::GOT-1.2*) |
|  | ZM8631 | - | *hpIs484* | pJH3437 (*Prgef-1::GOT-1::GFP*) |
|  | ZM9008 | - | *hp881* | pJH3621 (*targeting construct for*<br>*generating C-terminal GFP insertion at*<br>*the endogenous locus of GOT-1.2*) |
|  | ZM9077 | *ubr-1(hp684)* |  |  |
|  | ZM9084 | *ubr-1(hp821hp833)* |  |  |

## References

1. Pickart CM, Eddins MJ. Ubiquitin: structures, functions, mechanisms. Biochim Biophys Acta. 2004;1695(1-3):55–72.

2. Schwarz LA, Patrick GN. Ubiquitin-dependent endocytosis, trafficking and turnover of neuronal membrane proteins. Mol Cell Neurosci. 2012;49(3):387–93.

3. Ciechanover A, Iwai K. The ubiquitin system: from basic mechanisms to the patient bed. IUBMB Life. 2004;56(4):193–201.

4. Ciechanover A, Orian A, Schwartz AL. Ubiquitin-mediated proteolysis: biological regulation via destruction. Bioessays. 2000;22(5):442–51.

5. Hershko A, Ciechanover A. The ubiquitin system. Annu Rev Biochem. 1998;67:425–79.

6. Varshavsky A. The N-end rule: functions, mysteries, uses. Proc Natl Acad Sci U S A. 1996;93(22):12142–9.

7. Sriram SM, Kim BY, Kwon YT. The N-end rule pathway: emerging functions and molecular principles of substrate recognition. Nat Rev Mol Cell Biol. 2011;12(11):735–47.

8. Rao H, Uhlmann F, Nasmyth K, Varshavsky A. Degradation of a cohesin subunit by the N-end rule pathway is essential for chromosome stability. Nature. 2001;410(6831):955–9.

9. Stolz A, Besser S, Hottmann H, Wolf DH. Previously unknown role for the ubiquitin ligase Ubr1 in endoplasmic reticulum-associated protein degradation. Proc Natl Acad Sci U S A. 2013;110(38):15271–6.

10. Weaver BP, Weaver YM, Mitani S, Han M. Coupled Caspase and N-End Rule Ligase Activities Allow Recognition and Degradation of Pluripotency Factor LIN-28 during Non-Apoptotic Development. Dev Cell. 2017;41(6):665–73 e6.

11. Piatkov KI, Brower CS, Varshavsky A. The N-end rule pathway counteracts cell death by destroying proapoptotic protein fragments. Proc Natl Acad Sci U S A. 2012;109(27):E1839–47.

12. An JY, Seo JW, Tasaki T, Lee MJ, Varshavsky A, Kwon YT. Impaired neurogenesis and cardiovascular development in mice lacking the E3 ubiquitin ligases UBR1 and UBR2 of the N-end rule pathway. Proc Natl Acad Sci U S A. 2006;103(16):6212–7.

13. Zenker M, Mayerle J, Lerch MM, Tagariello A, Zerres K, Durie PR, et al. Deficiency of UBR1, a ubiquitin ligase of the N-end rule pathway, causes pancreatic dysfunction, malformations and mental retardation (Johanson-Blizzard syndrome). Nat Genet. 2005;37(12):1345–50.

14. Hwang CS, Sukalo M, Batygin O, Addor MC, Brunner H, Aytes AP, et al. Ubiquitin ligases of the N-end rule pathway: assessment of mutations in UBR1 that cause the Johanson-Blizzard syndrome. PLoS One. 2011;6(9):e24925.

15. Ottersen OP, Storm-Mathisen J. Glutamate- and GABA-containing neurons in the mouse and rat brain, as demonstrated with a new immunocytochemical technique. J Comp Neurol. 1984;229(3):374–92.

16. Mattson MP. Glutamate and neurotrophic factors in neuronal plasticity and disease. Ann N Y Acad Sci. 2008;1144:97–112.

17. DiAntonio A. Glutamate receptors at the Drosophila neuromuscular junction. Int Rev Neurobiol. 2006;75:165–79.

18. Brockie PJ, Maricq AV. Ionotropic glutamate receptors in Caenorhabditis elegans. Neurosignals. 2003;12(3):108–25.

19. Dong XX, Wang Y, Qin ZH. Molecular mechanisms of excitotoxicity and their relevance to pathogenesis of neurodegenerative diseases. Acta Pharmacol Sin. 2009;30(4):379–87.

20. Siegel S, Sanacora G. The roles of glutamate receptors across major neurological and psychiatric disorders. Pharmacol Biochem Behav. 2012;100(4):653–5.

21. Tasaki T, Mulder LC, Iwamatsu A, Lee MJ, Davydov IV, Varshavsky A, et al. A family of mammalian E3 ubiquitin ligases that contain the UBR box motif and recognize N-degrons. Mol Cell Biol. 2005;25(16):7120–36.

22. Xie Y, Varshavsky A. The E2-E3 interaction in the N-end rule pathway: the RING-H2 finger of E3 is required for the synthesis of multiubiquitin chain. EMBO J. 1999;18(23):6832–44.

23. Berndsen CE, Wolberger C. New insights into ubiquitin E3 ligase mechanism. Nat Struct Mol Biol. 2014;21(4):301–7.

24. Budhidarmo R, Nakatani Y, Day CL. RINGs hold the key to ubiquitin transfer. Trends Biochem Sci. 2012;37(2):58–65.

25. Hatakeyama S, Yada M, Matsumoto M, Ishida N, Nakayama KI. U box proteins as a new family of ubiquitin-protein ligases. J Biol Chem. 2001;276(35):33111–20.

26. Friedland AE, Tzur YB, Esvelt KM, Colaiacovo MP, Church GM, Calarco JA. Heritable genome editing in C. elegans via a CRISPR-Cas9 system. Nat Methods. 2013;10(8):741–3.

27. Sukalo M, Fiedler A, Guzman C, Spranger S, Addor MC, McHeik JN, et al. Mutations in the human UBR1 gene and the associated phenotypic spectrum. Human Mutation. 2014;35(5):521–31.

28. Brockie PJ, Mellem JE, Hills T, Madsen DM, Maricq AV. The C. elegans glutamate receptor subunit NMR-1 is required for slow NMDA-activated currents that regulate reversal frequency during locomotion. Neuron. 2001;31(4):617–30.

29. Maricq AV, Peckol E, Driscoll M, Bargmann CI. Mechanosensory signalling in C. elegans mediated by the GLR-1 glutamate receptor. Nature. 1995;378(6552):78–81.

30. Brockie PJ, Madsen DM, Zheng Y, Mellem J, Maricq AV. Differential expression of glutamate receptor subunits in the nervous system of Caenorhabditis elegans and their regulation by the homeodomain protein UNC-42. J Neurosci. 2001;21(5):1510–22.

31. Chalfie M, Sulston JE, White JG, Southgate E, Thomson JN, Brenner S. The neural circuit for touch sensitivity in Caenorhabditis elegans. J Neurosci. 1985;5(4):956–64.

32. Kawano T, Po MD, Gao S, Leung G, Ryu WS, Zhen M. An imbalancing act: gap junctions reduce the backward motor circuit activity to bias C. elegans for forward locomotion. Neuron. 2011;72(4):572–86.

33. Wicks SR, Roehrig CJ, Rankin CH. A dynamic network simulation of the nematode tap withdrawal circuit: predictions concerning synaptic function using behavioral criteria. J Neurosci. 1996;16(12):4017–31.

34. White JG, Southgate E, Thomson JN, Brenner S. The structure of the ventral nerve cord of Caenorhabditis elegans. Philos Trans R Soc Lond B Biol Sci. 1976;275(938):327–48.

35. Gray JM, Hill JJ, Bargmann CI. A circuit for navigation in Caenorhabditis elegans. Proc Natl Acad Sci U S A. 2005;102(9):3184–91.

36. Alkema MJ, Hunter-Ensor M, Ringstad N, Horvitz HR. Tyramine Functions independently of octopamine in the Caenorhabditis elegans nervous system. Neuron. 2005;46(2):247–60.

37. Faumont S, Rondeau G, Thiele TR, Lawton KJ, McCormick KE, Sottile M, et al. An image-free opto-mechanical system for creating virtual environments and imaging neuronal activity in freely moving Caenorhabditis elegans. PLoS One. 2011;6(9):e24666.

38. Ben Arous J, Tanizawa Y, Rabinowitch I, Chatenay D, Schafer WR. Automated imaging of neuronal activity in freely behaving Caenorhabditis elegans. J Neurosci Methods. 2010;187(2):229–34.

39. Chronis N, Zimmer M, Bargmann CI. Microfluidics for in vivo imaging of neuronal and behavioral activity in Caenorhabditis elegans. Nat Methods. 2007;4(9):727–31.

40. Piggott BJ, Liu J, Feng Z, Wescott SA, Xu XZ. The neural circuits and synaptic mechanisms underlying motor initiation in C. elegans. Cell. 2011;147(4):922–33.

41. Zheng Y, Brockie PJ, Mellem JE, Madsen DM, Maricq AV. Neuronal control of locomotion in C. elegans is modified by a dominant mutation in the GLR-1 ionotropic glutamate receptor. Neuron. 1999;24(2):347–61.

42. Gao S, Xie L, Kawano T, Po MD, Guan S, Zhen M. The NCA sodium leak channel is required for persistent motor circuit activity that sustains locomotion. Nat Commun. 2015;6:6323.

43. Liu P, Chen B, Mailler R, Wang ZW. Antidromic-rectifying gap junctions amplify chemical transmission at functionally mixed electrical-chemical synapses. Nat Commun. 2017;8:14818.

44. Haspel G, O’Donovan MJ, Hart AC. Motoneurons dedicated to either forward or backward locomotion in the nematode Caenorhabditis elegans. J Neurosci. 2010;30(33):11151–6.

45. White JG, Southgate E, Thomson JN, Brenner S. The structure of the nervous system of the nematode Caenorhabditis elegans. Philos Trans R Soc Lond B Biol Sci. 1986;314(1165):1–340.

46. Lim MA, Chitturi J, Laskova V, Meng J, Findeis D, Wiekenberg A, et al. Neuroendocrine modulation sustains the C. elegans forward motor state. Elife. 2016;5.

47. Gao S, Guan SA, Fouad AD, Meng J, Kawano T, Huang Y-C, et al. Excitatory Motor Neurons are Local Oscillators for Reverse Locomotion. Elife. 2017;in press.

48. Hayashi M, Yamada H, Uehara S, Morimoto R, Muroyama A, Yatsushiro S, et al. Secretory granule-mediated co-secretion of L-glutamate and glucagon triggers glutamatergic signal transmission in islets of Langerhans. J Biol Chem. 2003;278(3):1966–74.

49. Hirotsu K, Goto M, Okamoto A, Miyahara I. Dual substrate recognition of aminotransferases. Chem Rec. 2005;5(3):160–72.

50. Falk MJ, Zhang Z, Rosenjack JR, Nissim I, Daikhin E, Nissim I, et al. Metabolic pathway profiling of mitochondrial respiratory chain mutants in C. elegans. Mol Genet Metab. 2008;93(4):388–97.

51. Son J, Lyssiotis CA, Ying H, Wang X, Hua S, Ligorio M, et al. Glutamine supports pancreatic cancer growth through a KRAS-regulated metabolic pathway. Nature. 2013;496(7443):101–5.

52. Serrano-Saiz E, Poole RJ, Felton T, Zhang F, De La Cruz ED, Hobert O. Modular control of glutamatergic neuronal identity in C. elegans by distinct homeodomain proteins. Cell. 2013;155(3):659–73.

53. Kelly A, Stanley CA. Disorders of glutamate metabolism. Ment Retard Dev Disabil Res Rev. 2001;7(4):287–95.

54. Schild LC, Zbinden L, Bell HW, Yu YV, Sengupta P, Goodman MB, et al. The balance between cytoplasmic and nuclear CaM kinase-1 signaling controls the operating range of noxious heat avoidance. Neuron. 2014;84(5):983–96.

55. Starich TA, Xu J, Skerrett IM, Nicholson BJ, Shaw JE. Interactions between innexins UNC-7 and UNC-9 mediate electrical synapse specificity in the Caenorhabditis elegans locomotory nervous system. Neural Dev. 2009;4:16.

56. Hawkins RA. The blood-brain barrier and glutamate. Am J Clin Nutr. 2009;90(3):867S–74S.

57. Palmada M, Centelles JJ. Excitatory amino acid neurotransmission. Pathways for metabolism, storage and reuptake of glutamate in brain. Front Biosci. 1998;3:d701–18.

58. McKenna MC. The glutamate-glutamine cycle is not stoichiometric: fates of glutamate in brain. J Neurosci Res. 2007;85(15):3347–58.

59. Takeda K, Ishida A, Takahashi K, Ueda T. Synaptic vesicles are capable of synthesizing the VGLUT substrate glutamate from alpha-ketoglutarate for vesicular loading. J Neurochem. 2012;121(2):184–96.

60. Hobert O. The neuronal genome of Caenorhabditis elegans. WormBook. 2013:1–106.

61. Takeda K, Yanagida M. Regulation of nuclear proteasome by Rhp6/Ubc2 through ubiquitination and destruction of the sensor and anchor Cut8. Cell. 2005;122(3):393–405.

62. Turner GC, Du F, Varshavsky A. Peptides accelerate their uptake by activating a ubiquitin-dependent proteolytic pathway. Nature. 2000;405(6786):579–83.

63. Kitamura K, Katayama S, Dhut S, Sato M, Watanabe Y, Yamamoto M, et al. Phosphorylation of Mei2 and Ste11 by Pat1 kinase inhibits sexual differentiation via ubiquitin proteolysis and 14-3-3 protein in fission yeast. Dev Cell. 2001;1(3):389–99.

64. Sasaki T, Kojima H, Kishimoto R, Ikeda A, Kunimoto H, Nakajima K. Spatiotemporal regulation of c-Fos by ERK5 and the E3 ubiquitin ligase UBR1, and its biological role. Mol Cell. 2006;24(1):63–75.

65. Moss EG, Lee RC, Ambros V. The cold shock domain protein LIN-28 controls developmental timing in C. elegans and is regulated by the lin-4 RNA. Cell. 1997;88(5):637–46.

66. Humphrey SJ, James DE, Mann M. Protein Phosphorylation: A Major Switch Mechanism for Metabolic Regulation. Trends Endocrinol Metab. 2015;26(12):676–87.

67. Oliveira AP, Ludwig C, Picotti P, Kogadeeva M, Aebersold R, Sauer U. Regulation of yeast central metabolism by enzyme phosphorylation. Mol Syst Biol. 2012;8:623.

68. Fallahi GH, Sabbaghian M, Khalili M, Parvaneh N, Zenker M, Rezaei N. Novel UBR1 gene mutation in a patient with typical phenotype of Johanson-Blizzard syndrome. Eur J Pediatr. 2011;170(2):233–5.

69. Gammelsaeter R, Coppola T, Marcaggi P, Storm-Mathisen J, Chaudhry FA, Attwell D, et al. A role for glutamate transporters in the regulation of insulin secretion. PLoS One. 2011;6(8):e22960.

70. Jenstad M, Chaudhry FA. The Amino Acid Transporters of the Glutamate/GABA-Glutamine Cycle and Their Impact on Insulin and Glucagon Secretion. Front Endocrinol (Lausanne). 2013;4:199.

71. Davis MW, Hammarlund M. Single-nucleotide polymorphism mapping. Methods Mol Biol. 2006;351:75–92.

72. Doitsidou M, Poole RJ, Sarin S, Bigelow H, Hobert O. C. elegans mutant identification with a one-step whole-genome-sequencing and SNP mapping strategy. PLoS One. 2010;5(11):e15435.

73. Mello CC, Kramer JM, Stinchcomb D, Ambros V. Efficient gene transfer in C.elegans: extrachromosomal maintenance and integration of transforming sequences. Embo j. 1991;10(12):3959–70.

74. Frokjaer-Jensen C, Davis MW, Hopkins CE, Newman BJ, Thummel JM, Olesen SP, et al. Single-copy insertion of transgenes in Caenorhabditis elegans. Nat Genet. 2008;40(11):1375–83.

75. Tzur YB, Friedland AE, Nadarajan S, Church GM, Calarco JA, Colaiacovo MP. Heritable custom genomic modifications in Caenorhabditis elegans via a CRISPR-Cas9 system. Genetics. 2013;195(3):1181–5.

76. Hilliard MA, Bargmann CI, Bazzicalupo P. C. elegans responds to chemical repellents by integrating sensory inputs from the head and the tail. Curr Biol. 2002;12(9):730–4.

77. Abdel Rahman AM, Ryczko M, Pawling J, Dennis JW. Probing the hexosamine biosynthetic pathway in human tumor cells by multitargeted tandem mass spectrometry. ACS Chem Biol. 2013;8(9):2053–62.

78. Hung W, Hwang C, Po MD, Zhen M. Neuronal polarity is regulated by a direct interaction between a scaffolding protein, Neurabin, and a presynaptic SAD-1 kinase in Caenorhabditis elegans. Development. 2007;134(2):237–49.

79. Haspel G, O’Donovan MJ. A perimotor framework reveals functional segmentation in the motoneuronal network controlling locomotion in Caenorhabditis elegans. J Neurosci. 2011;31(41):14611–23.

